# Building bridges: the hippocampus constructs narrative memories across distant events

**DOI:** 10.1101/2020.12.11.422162

**Authors:** Brendan I. Cohn-Sheehy, Angelique I. Delarazan, Zachariah M. Reagh, Jordan E. Crivelli-Decker, Kamin Kim, Alexander J. Barnett, Jeffrey M. Zacks, Charan Ranganath

## Abstract

Life’s events are scattered throughout time, yet we often recall different events in the context of an integrated narrative. Prior research suggests that the hippocampus, which supports memory for past events, can sometimes support integration of overlapping associations or separate events, but the conditions which lead to hippocampus-dependent memory integration are unclear. We used functional brain imaging to test whether the ability to form a larger narrative (narrative coherence) drives hippocampal memory integration. During encoding of fictional stories, the hippocampus supported patterns of activity, including activity at boundaries between events, which were more similar between distant events that formed one coherent narrative, compared with events taken from unrelated narratives. Twenty-four hours later, the hippocampus preferentially supported detailed recall of coherent narrative events, through reinstatement of hippocampal activity patterns from encoding. These findings reveal a key function of the hippocampus: the dynamic integration of events into a narrative structure for memory.

## Introduction

It is well-established that although experience unfolds over a continuous timeline, memory for past experiences, or episodic memory (Tulving, 1972), can be segmented into individual units of time called “events” (Zacks, 2020). For example, say you are in the middle of reading a scientific article when you receive a phone call from your friend Sandra. Sandra proceeds to vent about an upcoming date, with a man who might be very charming or, alternatively, quite boorish. If you recall Sandra’s call later on, you would find it difficult to also recall your time reading the scientific article—information about two adjacent events can become segregated in memory (Clewett et al., 2020; Ezzyat & Davachi, 2011; Horner et al., 2016; Radvansky & Copeland, 2006; Swallow et al., 2011; Zacks et al., 2007). However, event segmentation may not provide the only organization for memory. For example, you might subsequently encounter Sandra at a café, slapping an unnamed man as she walks out. Although Sandra appears in temporally separated events, they coalesce to form a *narrative*: a larger unit of information which encompasses multiple events, and in which one’s understanding of each event is dependent on information from other events (Bartlett, 1932; Cohn-Sheehy et al., 2021; Graesser et al., 1994; Trabasso et al., 1984; van Dijk & Kintsch, 1983; Willems et al., 2020).

The ability to organize events into a narrative can be beneficial for memory (Bransford & Johnson, 1972; Kintsch, 1992; Rumelhart, 1977; Thorndyke, 1977; Trabasso et al., 1984). For example we recently demonstrated that, when temporally separated events could be assimilated into a narrative, they were recalled in greater detail than overlapping events involving the same character in different narratives (Cohn-Sheehy et al., 2021). Furthermore, narratives are more than the semantic information conveyed in words and sentences alone (Graesser et al., 1994; Kintsch, 1992; Trabasso et al., 1984; van Dijk & Kintsch, 1983). Although some models have suggested that semantic relatedness can determine whether words or sentences become integrated in memory (e.g. Reder & Anderson, 1980), we found that the memory advantage for coherent narrative events was over and above any advantage that could be observed at the word or sentence level (Cohn-Sheehy et al., 2021). Our findings suggested that episodic memory might be organized at a level above and beyond individual words, sentences, or events, on the basis of *narrative coherence*: the degree to which individual units of information can be interrelated within a single narrative representation (Bartlett, 1932; Graesser et al., 1994). In the present study, we tested the possibility that narrative coherence shapes the way the human brain encodes and retrieves events in memory.

Little is known about how the brain assembles narratives from individual events, but there is reason to think that the hippocampus plays a critical role. Many findings support the general idea that the hippocampus can build memories that integrate information across overlapping experiences (Horner et al., 2015; Kuhl et al., 2010; Schlichting et al., 2014, 2015; Shohamy & Wagner, 2008; Zeithamova & Preston, 2010; Zeithamova et al., 2012), although this might not depend on narrative coherence. Several studies have suggested that when one encounters information that overlaps with a previous event (e.g. a recurring “B” item in separated “A-B” and “B-C” pairs), this overlap can trigger the hippocampus to form associations between temporally distant experiences (Schlichting et al., 2014, 2015; Zeithamova & Preston, 2010; Zeithamova et al., 2012). Some evidence supports a potential mechanism for memory integration during encoding: encountering a new event might trigger the hippocampus to reinstate activity patterns which correspond with a prior overlapping event, such that information from the prior event becomes embedded into memory for the new event (Horner et al., 2015; Stawarczyk et al., 2020; Zeithamova et al., 2012). If the hippocampus can combine information across events during memory encoding, this might make it easier to recall both events later on.

However, overlapping information does not always result in memory integration—in fact, longstanding evidence suggests that when events share overlapping features (e.g. a recurring character), these events are more difficult to remember (Radvansky, 2012; Radvansky & Zacks, 1991). That is, overlapping events can compete with each other during memory retrieval (Radvansky, 2012; Radvansky & Zacks, 1991). Furthermore, some findings suggest that the hippocampus might hyper-differentiate between overlapping events (Chanales et al., 2017; Favila et al., 2016), and results from another study suggest that the hippocampus might simultaneously support the ability to segregate and integrate events that share overlapping features (Schlichting et al., 2015). Recent models have attempted to describe the conditions under which the hippocampus supports memory integration versus differentiation (Chanales et al., 2017, 2019; Horner et al., 2015; Kumaran, 2012; Ritvo et al., 2019). These models generally focus on simple associations between items and do not address any role for narrative coherence—though, as our recent behavioral findings suggest (Cohn-Sheehy et al., 2021), the ability to form a narrative might determine whether or not events become integrated in memory.

Recent evidence suggests that narrative coherence modulates activity in the hippocampus (Collin et al., 2015; Milivojevic et al., 2015). For instance, Milivojevic et al. (2015) used functional magnetic resonance imaging (fMRI) to analyze patterns of hippocampal activity evoked by repeated pairs of temporally adjacent events. Although events were initially unrelated to each other, participants were instructed that some event pairs would form a narrative with a subsequently-presented, linker event—when events could become linked, hippocampal activity patterns became more similar between paired events (Milivojevic et al., 2015). This finding suggested that, after one initially encodes events, patterns of activity in the hippocampus might become modified to support information about a larger narrative. Related findings from Collin et al. (2015) suggested that hippocampal pattern overlap between adjacent events might only be observed in participants who consciously recognize links between events. However, overlapping activity patterns are not equivalent to memory integration per se. If similar activity patterns reflect that events are integrated in memory, we would also expect hippocampal activity patterns would play a beneficial role when memory is assessed (i.e. during recall of events). Critically, no study to date has demonstrated whether narrative coherence modulates hippocampus-dependent memory integration across temporally distant events.

An additional possibility is that the hippocampus might integrate events into a coherent narrative, but only at specific moments during experience. Several recent studies (Baldassano et al., 2017; Ben-Yakov & Dudai, 2011; Ben-Yakov & Henson, 2018; Reagh et al., 2020; Sinclair et al., 2020) have found that the hippocampus becomes particularly active when people perceive transitions between events, or *event boundaries* (Newtson, 1973; Zacks, 2020). Although the significance of boundary-evoked activation is unclear, it might indicate that the hippocampus supports retrieval of information from past events in order to inform encoding of a new event (Griffiths & Fuentemilla, 2020; Stawarczyk et al., 2020). For instance, running into your friend Sandra would not only mark the start of a new event, but might also lead you to retrieve Sandra’s previous event from memory and integrate it with the new event (Kintsch, 1992; Ross & Bradshaw, 1994; Stawarczyk et al., 2020; Zacks, 2020). In support of this idea, recent electroencephalography (EEG) findings suggested that, at an event boundary, neurophysiological activity recapitulates information about a preceding, temporally-adjacent event (Silva et al., 2019; Sols et al., 2017). On the other hand, event boundaries provide a basis for segmenting events in memory (Newtson, 1973; Zacks, 2020). In other words, event boundaries might act as “walls” between temporally adjacent events—or, they might also serve as optimal points for the hippocampus to build “bridges” between temporally distant events.

Building on prior findings showing that narratives modulate activity in the hippocampus (Collin et al., 2015; Milivojevic et al., 2015, 2016), we sought to determine whether the hippocampus supports memory integration for temporally distant events that form a larger, coherent narrative, by examining hippocampal activity patterns during encoding and subsequent recall of realistic events. Additionally, we tested whether hippocampal activity patterns at event boundaries support integration across events. Functional magnetic resonance imaging (fMRI) was used to image brain activity while participants listened to fictional stories (**Fig. 1A**), in which recurring characters appeared in pairs of temporally separated events. Critically, these stories were written to include pairs of events that could form one Coherent Narrative, and pairs involving an overlapping character in Unrelated Narratives (see **Fig. 1B** for examples). These temporally separated events were embedded in longer stories which did not involve these recurring characters. For example, one four-minute story (Story 2 in **Fig. 1A**) followed a protagonist who, by chance, encountered the characters Sandra and Johnny, and in a subsequent story (Story 4 in **Fig. 1A**), a different protagonist also encountered Sandra and Johnny. In one version of these stories, the two events involving Sandra could be integrated into a Coherent Narrative (e.g., two events that describe a single dating experience) whereas the events involving Johnny were parts of Unrelated Narratives (e.g., searching for a lost recipe in Event 1, evading financial troubles in Event 2). However, both Coherent and Unrelated Narrative events were meaningfully distinct from the surrounding plot, such that events involving recurring characters could only form a larger narrative in the Coherent Narrative condition. Importantly, different versions of the stimuli were constructed and pseudo-randomly assigned to different subjects (see **Methods**), in order to minimize content-driven differences between Coherent and Unrelated Narrative events. Participants returned one day later, and they were scanned during detailed recall of the events involving each recurring character (**Fig. 1C**).

**Fig. 1:**
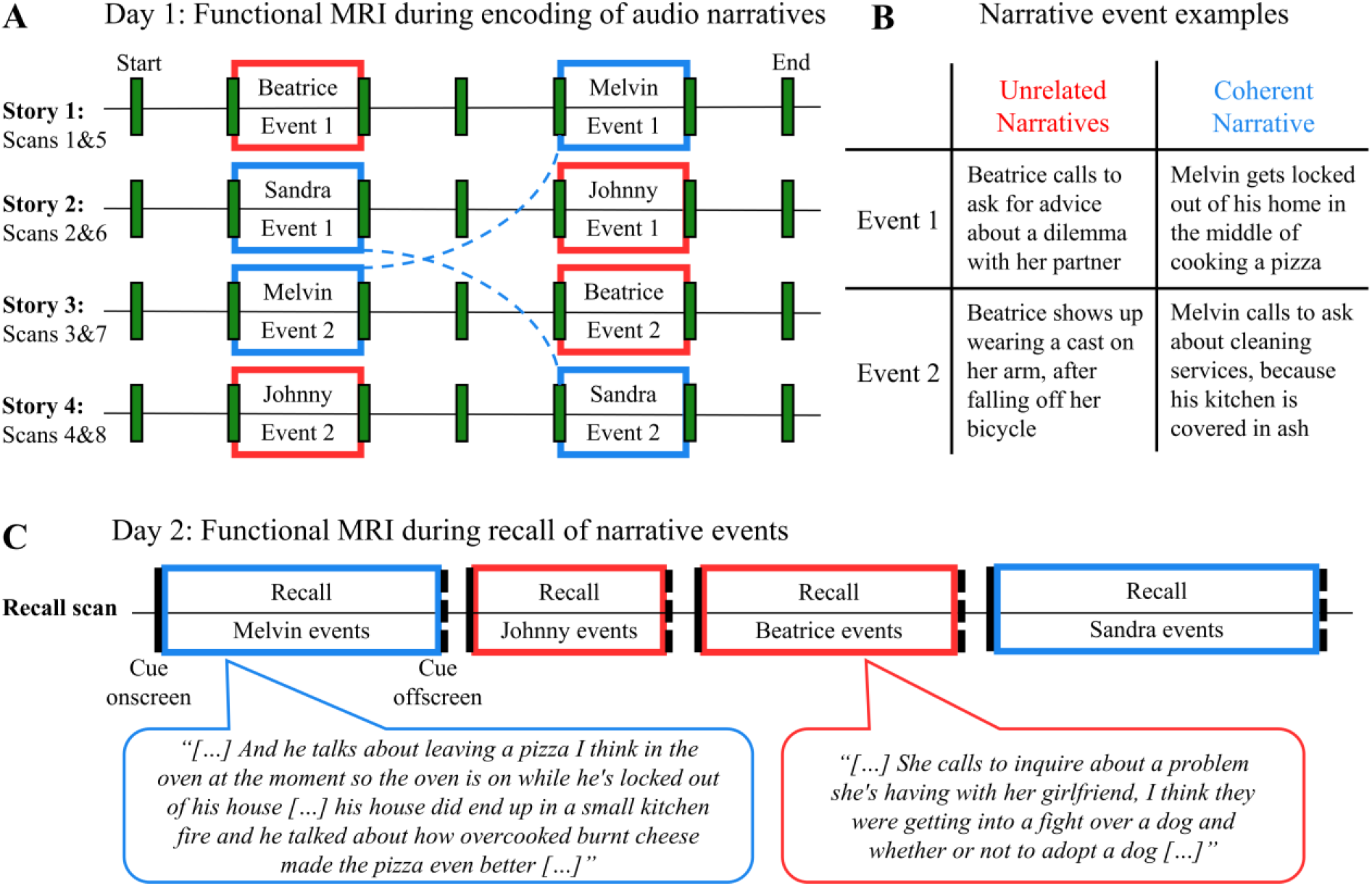
Experimental paradigm. **(A)** Narrative stimuli presented during functional MRI: On Day 1, four fictional stories were presented serially during scanning, as audio clips (240s each), which were punctuated by discrete event boundaries (green bars). Events involving recurring characters (Beatrice, Melvin, Sandra, Johnny; 40s each) did not relate to the more continuous plot events surrounding them. For two of these recurring characters (Blue boxes and dashed lines), two temporally-distant events could form one Coherent Narrative (e.g. Sandra Events 1 and 2). In contrast, for two other recurring characters (Red boxes), Events 1 and 2 belonged to Unrelated Narratives (e.g. Johnny). For each participant, side-characters were randomly assigned to either the Coherent Narrative or Unrelated Narratives conditions (i.e. this is one example; see Methods). **(B)** Examples of narrative events: Synopses are provided for possible pairs of Coherent Narrative and Unrelated Narrative events. **(C)** Recall assessment during functional MRI: During scanning on Day 2, participants were asked to recall events involving each recurring character from the stories presented on Day 1 (Coherent Narratives = Blue; Unrelated Narratives = Red). The amount of time spent on recall, and the amount of details recalled, varied from character to character. Recall excerpts are presented within speech bubbles.

We hypothesized that the hippocampus would play a disproportionate role in supporting memory for Coherent Narrative events, as compared with Unrelated Narrative events. We tested this prediction by determining whether hippocampal activity patterns during memory encoding carried shared information across separate events that form a coherent narrative, and whether activity patterns during these time periods were reinstated during recall of these events, as compared to reinstatement during recall of Unrelated Narrative events.

## Results

### Hippocampal activity bridges coherent narrative events during memory encoding

As described above, our main hypothesis was that hippocampal activity might bridge distant events, if the events can form a coherent narrative. To assess this possibility, we identified the unique pattern of hippocampal activity (voxel pattern) corresponding to each story event (see **Methods**; **Fig. 2A**), and investigated the degree to which voxel patterns were shared between events which formed a Coherent Narrative (CN) versus events which were taken from Unrelated Narratives (UN). We hypothesized that narrative coherence would determine whether the hippocampus could integrate temporally-distant events, and if so, we would expect higher pattern similarity between CN than UN events. Alternatively, the hippocampus might support the integration of any overlapping events (i.e. via a shared character), and if so, we would not expect any difference in pattern similarity between CN and UN events.

**Fig. 2:**
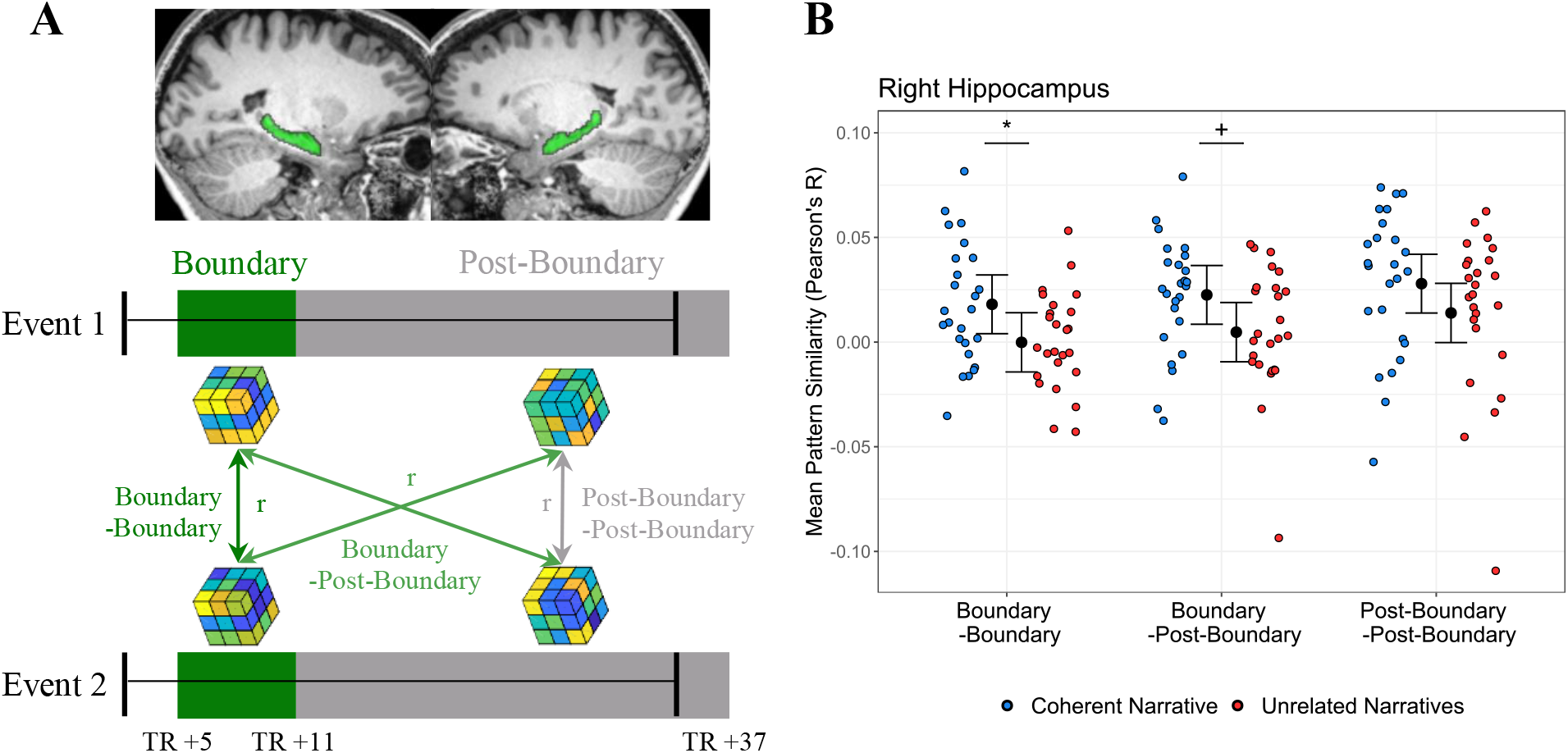
Right hippocampus activity patterns bridge events that form a coherent narrative. **(A)** Across-event pattern similarity analysis: within the left and right hippocampus (plotted on an individual subject’s structural MRI, in green),the unique spatial pattern of activity (“voxel pattern”, represented by cubes) was modeled for Boundary (dark green rectangles, TRs +5 to +11) and Post-Boundary epochs (grey rectangles, TRs +12 to +37). Black bars represent the real-time beginning and ending of each Coherent Narrative and Unrelated Narrative event during scanning—epoch definitions account for the lagged BOLD response. Pattern similarity (Pearson’s r) was calculated between voxel patterns at Event 1 and Event 2, and selectively averaged to yield three epoch-by-epoch measures of similarity: Boundary-Boundary (dark green line), Boundary-Post-Boundary (light green lines), and Post-Boundary-Post-Boundary (grey line). **(B)** Right hippocampus pattern similarity: Pearson’s correlations between Event 1 and 2 epochs were computed for Coherent Narrative and Unrelated Narrative events. Mean pattern similarity for individual subjects (colored dots) is plotted on the basis of which epoch was being examined, and whether events could form Coherent Narratives or not (Unrelated Narratives). Mixed model fits predicting pattern similarity are visualized as 95% confidence intervals (see text for model details). Key: Blue=Coherent Narratives, Red=Unrelated Narratives, +=p<0.10, *=p<0.05.

There was also reason to suspect that activity during event boundaries—which reliably engage the hippocampus—would contribute to memory integration (e.g. Griffiths & Fuentemilla, 2020). Each story was written to include *a priori* event boundaries (green bars in **Fig. 1A**), which reliably evoked the perception of event boundaries in an independent sample (point-biserial correlation: r=0.85, p<0.0001; **Fig. S1A**; see **Methods**). As in previous studies (e.g. Baldassano et al., 2017; Ben-Yakov & Henson, 2018; Reagh et al., 2020), *a priori* event boundaries evoked increased activation in the right and left hippocampus (**Fig. S1B**). The period of time which corresponded to the onset and duration of significant boundary-evoked activation in the hippocampus (TRs 5-11, ∼6.1-14.64 seconds following the onset of an event; see **Fig. S1B**) is subsequently referred to as the Boundary epoch (**Fig. 2A**). The Boundary epoch accounted for the lag between the real-time onset of an event, and an increase in activation on functional MRI (i.e. the hemodynamic response lag). For comparison, we also defined a Post-Boundary epoch (**Fig. 2A**), in order to examine patterns of activity during the remainder of an event. Because each event spanned 33 timepoints, the Post-Boundary epoch encompassed 26 timepoints following the Boundary epoch (TRs 12-37, ∼14.64-46.36 seconds following the onset of an event).

We computed and selectively averaged voxel pattern similarity between the Boundary and Post-Boundary epochs of paired events (**Fig. 2A**), and subsequently analyzed pattern similarity values within the right and left hippocampus using a multi-level mixed effects model (see **Methods**, Formula 1). This model tested whether Narrative Coherence [CN vs UN] would modulate pattern similarity across events, and whether any effects of Narrative Coherence would be specifically driven by similarity between Boundary epochs (i.e. an interaction between Narrative Coherence and Epoch). Furthermore, the mixed model accounted for nuisance regressors which might confound effects of interest: particular subjects, specific events which were presented to subjects, and overall activation at boundaries.

As shown in **Fig. 2B**, within the right hippocampus, this model (Akaike Information Criterion: AIC = -1039) revealed a significant effect of Narrative Coherence (*F*(1,11.33)=6.97, p=.02). This finding reflected that, even after accounting for potential confounds, pattern similarity in the right hippocampus was higher for CN events, compared with UN events (see **Fig. 2B** for model estimates). However, there was no significant interaction between Narrative Coherence and Epoch (*F*(2,257.71)=0.09, p=.92; ps≥0.10 for other fixed effects). Because the interaction was not significant, we cannot conclude that the Narrative Coherence effect was driven by the Boundary epoch alone. These findings suggested that during encoding of an event (either Boundary or Post-Boundary epochs), activity patterns in the right hippocampus might bridge distant events that form a coherent narrative. A follow-up analysis of timepoint-by-timepoint correlations between events revealed a similar pattern of findings (see **Fig. S2**).

This pattern of findings was not evident in the left hippocampus (**Figs. S2-3**), wherein the mixed model (AIC = -1103) revealed only a significant effect of overall boundary activation on pattern similarity (*F*(1,289.69)=12.18, p<.001; for other fixed effects, ps>.36). Interestingly, an exploratory analysis suggested that findings within the right hippocampus might be specific to the right posterior hippocampus, and not the right anterior hippocampus (**Fig. S4**); this analysis should be interpreted with caution.

Additionally, we sought to determine whether effects of narrative coherence were driven by semantic features of narrative text. Namely, if the hippocampus simply supports information about words and sentences (Duff et al., 2020; Piai et al., 2016; Solomon et al., 2019), it is possible that hippocampal pattern similarity might simply reflect the degree to which this kind of lower-level semantic information overlaps across events. We therefore used the Universal Sentence Encoder (Cer et al., 2018) to quantify textual similarity between events (see **Methods**), and we included textual similarity as an additional factor in mixed models of hippocampal pattern similarity (see **Methods,** Formula 2). Within the right hippocampus, this model (AIC = - 1037) revealed a significant effect of Narrative Coherence (*F*(1,12.06)=5.56, p=.036; other ps>0.14). This finding reflected that, even after accounting for lower-level semantic similarity between events, pattern similarity in the right hippocampus was higher for CN events, compared with UN events. This suggested that within the right hippocampus, the degree to which narrative coherence modulated pattern similarity across events was not purely driven by lower-level semantic overlap between events. This pattern of findings was not evident in the left hippocampus (AIC = -1101; effect of overall boundary activation, *F*(1,288.70)=12.17, p<.001; all other effects, ps>.58).

### Hippocampal activity that bridges coherent narrative events is reinstated during recall

The preceding findings supported the idea that, during memory encoding, patterns of activity in the right hippocampus are preferentially shared across distant events that form a coherent narrative. If activity patterns in the right hippocampus provided a basis for integrating Coherent Narrative events during memory encoding, we would expect that these activity patterns would play some role in subsequently retrieving these events from memory. Namely, we would expect that right hippocampal voxel patterns from encoding would be more highly reinstated during recall of CN events compared with UN events. Participants were scanned during recall of events involving CN and UN characters (**Fig. 1C**), providing an opportunity to compare voxel patterns between encoding and recall (i.e. “encoding-retrieval similarity”; **Fig. 3A**). Because preceding analyses revealed pattern similarity effects that spanned both Boundary and Post-Boundary epochs, we pooled timepoints from both epochs to measure reinstatement of whole-event activity patterns from encoding (**Fig. 3A**).

**Fig. 3:**
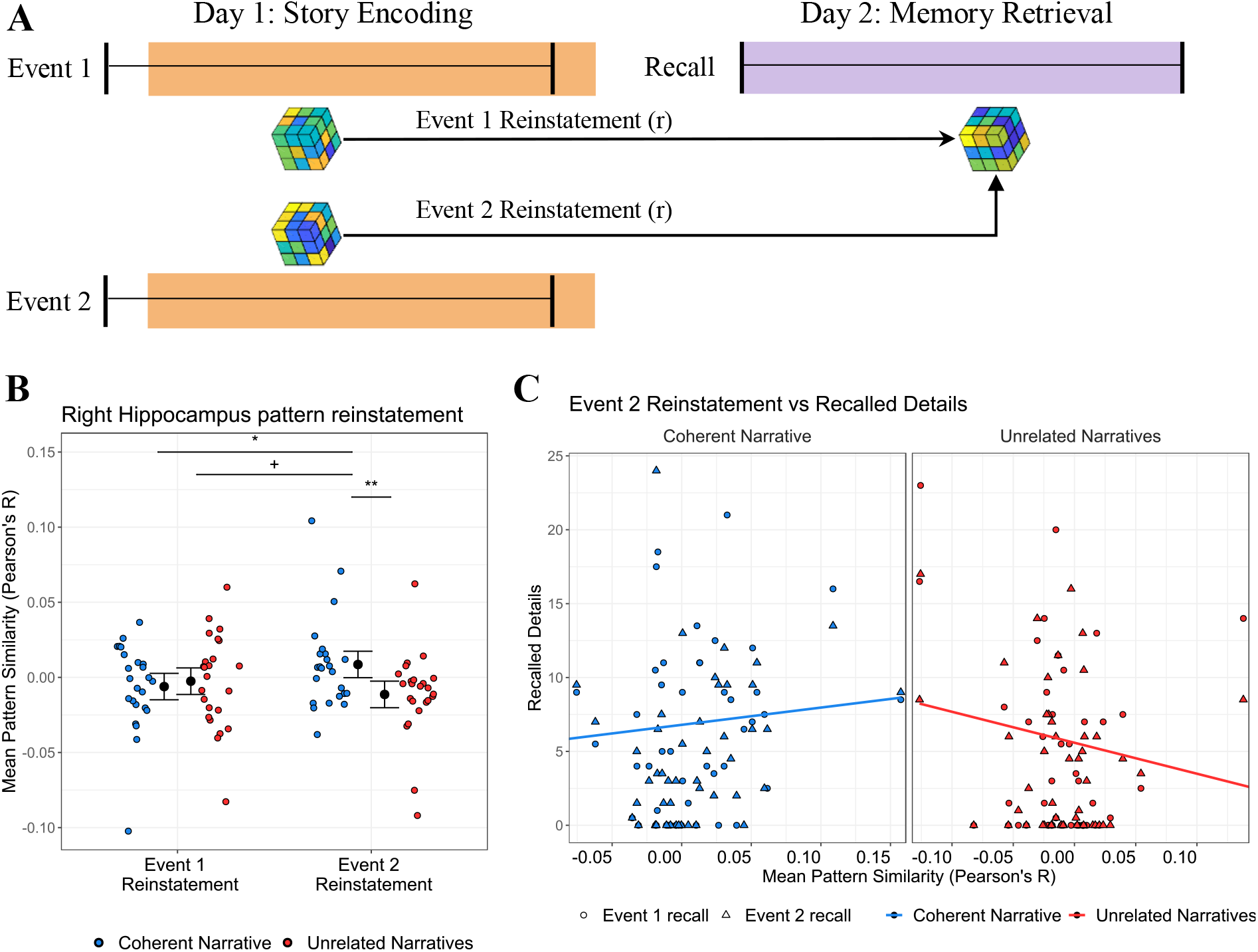
Reinstatement of hippocampal activity from memory encoding supports recall of Coherent Narrative events. **(A)** Encoding-retrieval similarity: Pattern similarity (Pearson’s r) was calculated between activity from each event involving a recurring character at encoding (Event 1 or 2), and activity during delayed recall of events involving that character. **(B)** Event 2 pattern reinstatement is higher for Coherent Narrative recall: Encoding-retrieval similarity for each subject (colored dots) is plotted for each epoch of each event during recall. 95% confidence intervals represent mixed model estimates (see text). **(C)** Event recall predicted by Event 2 pattern reinstatement: Recalled details from Event 1 (circles) and Event 2 (triangles) are plotted for each subject, predicted by Event 2 pattern reinstatement. Colored lines represent mixed model estimates (see text). Key: Blue=Coherent Narrative, Red=Unrelated Narratives, +=p<.10, *=p<.05, **=p<.01.

In order to accurately predict how reinstatement might work, it is important to consider how integration may have unfolded during encoding. During encoding, there were two events involving each CN and UN character (Event 1 and Event 2; see **Fig. 1A-B**). Because events were presented sequentially, Event 2 was the event which determined whether there was any relationship between the two events. If overlapping associations (e.g. a recurring character) drive the hippocampus to reinstate activity patterns across events (e.g. Zeithamova et al., 2012), then we might expect that hippocampal activity patterns during recall would reflect reinstatement of patterns evoked during encoding of Event 2 for both conditions. Alternatively, Event 2 might have served as a key period for integration specifically in the CN condition, in which case we would expect that reinstatement of information from Event 2 would be especially critical for subsequent recall of CN events. In contrast, we expected that it was unlikely that reinstatement of information from Event 2 would support recall of the two events in the UN condition, because these events could not be integrated into a larger narrative.

Based on this rationale, we hypothesized that right hippocampal activity patterns during encoding of Event 2 would be more highly reinstated during recall of CN events than UN events. To test this hypothesis, for each CN and UN character, we analyzed the degree to which right hippocampal voxel patterns during Event 2 encoding were reinstated during recall (see **Fig. 3A**, **Methods**). We also analyzed reinstatement of activity patterns from Event 1—however, because we suspected that Event 2 served as the critical period for integration, we did not expect any difference in reinstatement of activity from Event 1 between CN and UN events. To test our hypothesis, we performed a multi-level mixed effects model (see **Methods**, Formula 3) which tested whether encoding-retrieval similarity would be modulated by Narrative Coherence [CN vs UN events], and whether an effect of Narrative Coherence would be limited to reinstatement of Event 2 (i.e. an interaction between Narrative Coherence and Event Number). The model statistically accounted for several nuisance variables: the amount of time spent on recall (28-299 TRs, ∼ 0.5-6 minutes per recall cue), particular subjects, and particular event content.

This model (AIC = -1334.7) revealed a trending main effect of Narrative Coherence (*F*(1,303.98)=3.60, p=0.059), which was qualified by a significant interaction between Narrative Coherence and Event Number (*F*(1,343.46)=7.72, p=0.006); no other fixed effects were significant (ps>.48). As shown in **Fig. 3B**, model estimates revealed that the interaction was driven by an effect of higher reinstatement for CN than UN Events, and that this effect was specific to Event 2. An exploratory analysis revealed a similar pattern of findings when examining pattern reinstatement from Boundary epochs alone (**Fig. S5A-B**).

These findings suggested that, in the right hippocampus, patterns of activity from encoding of Event 2 were preferentially reinstated during recall of events which formed a Coherent Narrative. In line with our predictions, this suggested that Event 2 provided an opportunity for memory integration, and that hippocampal activity patterns during Event 2 may have integrated distant events which formed a coherent narrative.

### Narrative coherence modulates the relationship between hippocampal pattern reinstatement and detailed recall

Having established that information from Event 2 was preferentially reinstated during recall of CN events, our next analyses assessed whether this reinstatement effect was behaviorally relevant. In a previous behavioral study, we found that, across three experiments, participants were able to recall more details about CN than UN events (Cohn-Sheehy et al., 2021). In the present study, the number of details recalled from CN events (mean=10.71 details per cue, SD=9.4, max=41.5) was numerically higher than recall of details from UN events (mean=8.65 details per cue, SD=9.1, max=40.0), though this effect did not meet the threshold for statistical significance (*t*(24) =1.98, p=.059, Cohen’s d=0.22). Examination of the data revealed a large degree of inter-subject variance in recall performance, and we considered the possibility that this variance might be explained by inter-subject differences in hippocampal activity that supports recall. If so, we would expect that recall of CN events should be predicted by reinstatement of right hippocampus activity patterns during recall.

Following up on our analyses of encoding-retrieval similarity (**Fig. 3B**), we first tested whether reinstatement of right hippocampus activity patterns from CN Event 2 was predictive of how many details participants recalled about CN events. In line with our prediction, there was a significant positive correlation between mean CN Event 2 reinstatement and mean recalled details for CN events (*r*=0.41, p=0.048). That is, the degree to which Event 2 patterns were reinstated during recall predicted the degree to which participants recalled CN events in detail. For comparison, we also tested whether UN Event 2 reinstatement would predict detailed recall. In contrast, there was a negative, but non-significant, across-subjects correlation between mean UN Event 2 reinstatement and mean recall of UN events (*r*=-0.23, p=0.28). This initially suggested that narrative coherence might modulate the relationship between right hippocampal pattern reinstatement and detailed recall. We investigated this possibility further by conducting direct analyses with mixed-effects models.

As described above (**Fig. 3B**), evidence from analyses of encoding-retrieval similarity suggested that Event 2 provided an opportunity for memory integration during encoding, enabling CN events, and not UN events, to become integrated in memory. If so, then we might expect that reinstatement of right hippocampus activity patterns from CN Event 2 would not only predict how well Event 2 was recalled, but would also predict how well Event 1 was recalled, reflecting that these events became integrated in memory. In contrast, if UN events were not integrated in memory, we would not expect reinstatement of UN Event 2 activity patterns to predict detailed recall of both UN Events. Alternatively, if our pattern similarity findings did not reflect memory integration, we might expect that the right hippocampus would generally support recollection of individual events. If so, we might expect that reinstatement of activity from any event (i.e. either CN or UN Event 2) would predict only the number of details recalled about that specific event (i.e. recall of CN Event 2, recall of UN Event 2).

To assess whether and how memory integration occurred, we performed a multi-level mixed effects model (see **Methods**, Formula 4), to test whether and how recalled details would be predicted by Event 2 pattern reinstatement. Moreover, the model tested whether the relation between recall and pattern reinstatement would be modulated by Narrative Coherence [CN vs UN events]. The model also tested whether Event 2 pattern reinstatement would differentially predict the number of details recalled about Event 1 versus Event 2. After accounting for nuisance regressors (recall duration, individual subjects, specific events), this model (AIC=1120.5) revealed a significant main effect of Narrative Coherence (*F*(1,160.17)=4.40, p=0.037), which was qualified by a significant interaction of Narrative Coherence and Event 2 Reinstatement (*F*(1,156.42)=5.20, p=0.024). No other fixed effects were significant (all ps>.34), suggesting that Event 2 Reinstatement did not differentially predict recalled details for Event 1 versus Event 2, and that there were no overall differences in recall of Event 1 versus Event 2. A similar pattern of findings was revealed when excluding events with zero recalled details from this model (AIC=796.91; significant interaction of Narrative Coherence and Event 2 Reinstatement, *F*(1,99.01)=4.45, p=0.037).

As shown in **Fig. 3C**, the significant interaction reflected that in the right hippocampus, the relationship between Event 2 pattern reinstatement and recalled details (from either event) was modified by whether events could form a Coherent Narrative. To break down this interaction further, we performed a pairwise comparison of linear trends that were predicted by the mixed model (shown in **Fig. 3C**). This comparison revealed that the relationship between recalled details and Event 2 Reinstatement was significantly more positive for CN events than UN events (*t*(156)=2.28, p=0.024). Consistent with our hypothesis, this suggests that narrative coherence modulated the degree to which hippocampal pattern reinstatement from the second of two overlapping events predicted any benefit for subsequent recall. A similar pattern of findings was revealed when limiting the model to reinstatement from the Boundary epoch (**Fig. S5C**).

### Supplemental analyses of cortical networks

Our study was specifically designed to test *a priori* hypotheses about hippocampal contributions to memory. However, given the novelty of our design, it is possible to generate many testable hypotheses about any other brain areas that might be involved in this task. For instance, in the course of peer review, one reviewer requested that we test additional hypotheses about networks of cortical regions which interact with the hippocampus—namely, the posterior medial and anterior temporal subnetworks of the default mode network (Ranganath & Ritchey, 2012; Reagh & Ranganath, 2018). For completeness, we have included supplemental cortical analyses (**Figs. S6-7**). These analyses revealed significant effects of narrative coherence within the anterior temporal network and auditory cortex, but not within the posterior medial network.

## Discussion

In the current study, we investigated whether the hippocampus supports a narrative-level organization for episodic memories, by integrating temporally distant events into a larger narrative. In support of this hypothesis, activity patterns in the right hippocampus during memory encoding were more similar across two temporally separated events that formed a Coherent Narrative, than across Unrelated Narrative events with an overlapping character. Twenty-four hours later, during memory retrieval, activity patterns in the right hippocampus were more highly reinstated for Coherent Narratives than for Unrelated Narratives. In particular, activity reinstatement was disproportionately higher for the second of two events that formed a Coherent Narrative—that is, the event which provided the clearest opportunity to meaningfully integrate information across events. Furthermore, narrative coherence determined the degree to which reinstatement of hippocampal activity patterns from the second event predicted the ability to recall details about events.

Taken together, these findings suggest that, rather than simply integrating memories of overlapping experiences, the hippocampus supports the construction of narratives that encompass temporally distant events. A considerable body of evidence suggests that hippocampus-dependent memory processes can support memory integration—the ability to put together information across two simple associations with an overlapping element with a prior event (Kuhl et al., 2010; Schlichting et al., 2014, 2015; Zeithamova & Preston, 2010; Zeithamova et al., 2012)—though it is also the case that the hippocampus can assign highly differentiated representations to overlapping associations (Chanales et al., 2017; Favila et al., 2016). In related work, Koster et al. (2018) found evidence suggesting that the hippocampus keeps event representations distinct during memory encoding, instead supporting integration of overlapping associations on the fly during memory retrieval, consistent with a computational model (Kumaran, 2012).

Although important questions remain about dynamics of memory integration in these studies, it is important to clarify differences between the present study and previous work on memory integration. Studies of memory integration typically focus on overlapping associations that share the same element—for instance, two houses or scenes that are associated with the same face (e.g. Preston et al., 2004; Shohamy & Wagner, 2008). In the present study, both Coherent and Unrelated Narrative events shared overlapping characters, yet hippocampal pattern similarity was significantly higher across CN events, and CN event patterns were preferentially reinstated during recall 24 hours later. Thus, overlap across events, while potentially necessary, was not sufficient to integrate information across distant events. Furthermore, in contrast with prior studies, the present study investigated memory integration in the context of complex, realistic events and narratives.

A few studies have shown that hippocampal activity during encoding of temporally adjacent events can be sensitive to narrative coherence (Collin et al., 2015; Milivojevic et al., 2015). Building on these studies, the present findings directly support a relationship between narrative coherence effects in the hippocampus and detailed recall of temporally distant events. In other words, the present findings suggest that narrative coherence does more than modulate hippocampal activity across repeatedly presented events—rather, they directly support the hypothesis that narrative coherence drives hippocampal memory integration, and that this integration can even take place across events that are encountered at disparate times.

Beyond the present study and a few others (Collin et al., 2015; Milivojevic et al., 2015, 2016), most neuroimaging studies have not demonstrated significant hippocampal support for narrative construction. Many studies have demonstrated cortical support for the representation of narrative structure (e.g. Aly et al., 2018; Baldassano et al., 2017; Chang et al., 2021; Chen et al., 2017; Heusser et al., 2021). For instance, Aly et al. (2018) scanned participants while they repeatedly watched movie clips in which the order of events was intact, and other clips in which the order of events was scrambled so that it did not preserve the narrative structure. Results showed that activity within the posterior medial network (PMN)—a set of regions in the medial and ventrolateral parietal cortex—was more similar across repetitions of films with intact narratives, compared to repetitions of scrambled films, consistent with a role in narrative construction. This was not reported for the hippocampus, although functional connectivity between the PMN and hippocampus increased across repetitions of scrambled films (see also, Chen et al., 2016; Michelmann et al., 2020). In contrast to the results of Aly et al. (2018), supplemental analyses in our study (**Fig. S6**) did not reveal significant effects in the PMN.

Although this is beyond the scope of our findings, there are potentially a number of reasons why the hippocampus and PMN might play different roles in constructing memories for narratives. One important fact is that PMN activity patterns tend to be stable across the entirety of an event, and in fact, most fMRI studies of narrative structure have focused on characterization of pattern similarity across different individuals (e.g. Baldassano et al., 2017; Chen et al., 2017; Oedekoven et al., 2017; Reagh & Ranganath, 2021). In contrast, hippocampal activity tends to be more dynamic (e.g. Ben-Yakov & Henson, 2018). For instance, Reagh & Ranganath (2021) compared activity patterns across a 40-second movie, and results showed that PMN patterns remained highly correlated across the repetitions during the middle and end of each film. In contrast, hippocampal patterns primarily carried information about the time period following the onset of the film. This finding is interesting, because some of the key hippocampal findings in the present study were seen during the Boundary epoch.

To be clear, we cannot conclude, nor can we rule out, that event boundaries played a unique or disproportionate role in integrating temporally distant events in memory. Our hypothesis-driven analyses demonstrated that pattern similarity across CN events was higher than similarity across UN events during the Boundary epoch (i.e., onset of the event) and between the Boundary epoch and Post-Boundary epoch (**Fig. 2B**; see also **Fig. S2C**). In other words, hippocampal activity during the Boundary epoch carried similar information across Coherent Narrative events. Notably, several models suggest that when a new event begins (i.e. at an event boundary), people can update their current representation of an event, by retrieving information about relevant past events (Franklin et al., 2020; Griffiths & Fuentemilla, 2020; Lu et al., 2020; Stawarczyk et al., 2020; Zacks et al., 2007). In a recent model, Lu et al. (2020) demonstrated that it can be computationally advantageous to restrict hippocampal encoding and retrieval of cortical states to periods around an event boundary, and this idea fits with the established idea that people often rely on episodic retrieval to build mental representations of an ongoing event (Franklin et al., 2020; Griffiths & Fuentemilla, 2020; Ross & Bradshaw, 1994; Stawarczyk et al., 2020; Wahlheim & Jacoby, 2013; Zacks et al., 2007). In other words, even if event boundaries serve as a “wall” between temporally adjacent events, hippocampal retrieval at boundaries might support the construction of a “bridge” between temporally distant events.

Another key finding in the present study is that hippocampal activity patterns during CN events were preferentially reinstated during memory retrieval, and this reinstatement effect was more predictive of individual differences in memory for CN than UN events. In our previous behavioral study (Cohn-Sheehy et al., 2021), memory performance was significantly higher for events that formed a coherent narrative than for overlapping, but unrelated events. In order to adapt the paradigm for fMRI voxel pattern analyses, we adjusted this paradigm to include repeated presentation of each story (i.e. to increase the number of trials per condition). It is likely that repeated presentation enabled participants to encode more information about UN events, which would explain why detailed recall was numerically, but not significantly higher for CN than UN events. Nonetheless, participants showed preferential reinstatement of information from Event 2—precisely when integration would be expected to occur—during recall of CN events. If hippocampal activity simply reflected successful encoding of the second event, we would expect to see that reinstatement would be equally predictive of memory performance for CN and UN events, since memory performance was not significantly different across the two conditions. In fact, reinstatement of information during this key period when participants could build a coherent narrative was predictive of memory for those narratives, whereas reinstatement of information from an independent event was not significantly predictive of memory performance, and if anything, it appeared to be negatively correlated with behavior.

In closing, the current findings suggest that even though life’s events can occur at disparate times, the hippocampus can form memories that integrate these events into a larger, coherent narrative. By bridging the divide between distant events, the hippocampus may support a narrative-level architecture for episodic memory.

## Methods

### Participants

#### Functional MRI study

Thirty-six right-handed participants aged 18-32 years-old (mean age= 24.4, SD= 3.6; 24 female, 12 male) were recruited to participate in the imaging study. Among other criteria, participants were screened for native English language fluency. After informed consent was obtained, additional screening at the time of scanning revealed that two of these participants had contraindications for MRI scanning, and thus could not be included. Of the thirty-four remaining participants scanned on Day 1, seven participants were excluded because they did not return for the recall session on Day 2 (due to unanticipated issues with scanner or peripherals, or substantial motion or imaging artifacts on Day 1). For one additional subject, a scanner error precluded imaging during two of the eight story presentation scans on Day 1, however this participant was able to complete the story listening task (without concurrent scanning during the final two blocks), and was therefore retained for participation on Day 2. As a result, twenty-seven participants were scanned on Day 2. Two of these subjects were excluded because they were unable to successfully recall any details during the recall task. In all, twenty-five participants (mean age = 25.2, SD = 3.7; 15 female, 10 male) were included in the analyses reported in this paper, twenty-four of whom had sufficient data quality during recall to be included in encoding-retrieval similarity analyses.

#### Event boundary ratings

An independent sample of eighteen participants aged 18-26 years-old (mean age=20.4 , SD=2.0; 12 female), screened for native English language fluency, provided informed consent and were recruited to provide annotations of event boundaries (i.e. no imaging data was collected).

### Experimental Stimuli

The stimulus design is depicted in **Fig. 1A-B**, and has been described elsewhere (Cohn-Sheehy et al., 2021). We constructed four fictional stories in which we manipulated whether temporally-distant events could form a coherent narrative (**Data S1**). Story audio was scripted, recorded, and thoroughly edited by the first author (BICS) to ensure equivalent lengths for all sentences (5s each), events (8 sentences/40s each), and stories (6 events/240s each), and to avoid any differences in perceptual distinctiveness (i.e. similar audio quality and smooth transitions between sentences). Importantly, most 40s events were initiated and terminated by shifts in information that are known to trigger the perception of event boundaries (e.g. characters entering or exiting, temporal shifts like “20 minutes later”) (Ezzyat & Davachi, 2011; Zacks et al., 2009). Two additional boundaries marked the start and end of each story, respectively. These seven *a priori* event boundaries for each story (green bars in **Fig. 1A**; confirmed by human raters, **Fig. S1A**) provided the basis for subsequent fMRI analysis.

Two of the stories are centered on a character named Charles, who is attempting to get a big scoop for a newspaper, and two are centered on a character named Karen, who is attempting to find employment as a chef. To examine the effects of narrative-level organization on memory, we incorporated events involving key “side-characters” into each story (**Fig. 1A**). Two stories involved “Beatrice” and “Melvin” as side-characters, and two involved “Sandra” and “Johnny.” Each side-character appeared in two temporally-distant, distinct events that occurred 6-12 minutes apart (lags matched between CN and UN conditions), in disparate contexts across two unrelated stories. For instance, Sandra’s first event (Event 1) occurs in Story 2, when she calls Charles, who is sitting at a park at noon on a Tuesday. Sandra’s second event (Event 2) occurs in Story 4, when Karen runs into her at a French restaurant, at early evening the next day. Events involving the side-characters were tangential to adjacent main plot events which centered on Charles and Karen (i.e. they were “sideplots”).

Critically, for two recurring side-characters, these sideplot event pairs could form a Coherent Narrative (CN) about one particular situation involving that side-character. In contrast, for the other two recurring side-characters, each sideplot event described a different situation involving that side-character, such that the two sideplot events could not easily form a singular coherent narrative (Unrelated Narratives, UN). Each story contained one CN event and one UN event (see **Fig. 1A**). Because the stimulus design controlled for several other features that could support integration of sideplot events (temporal proximity, contextual similarity, attention to intervening main plot events), only CN events, and not UN events, could be easily integrated into a larger narrative.

We sought to control for any effects of specific event content or character identity that could confound the coherence manipulation, by randomizing CN and UN event content across subjects. For each side-character, we created two alternate pairs of CN events (e.g. Sandra Events 1 and 2, version A; Sandra Events 1 and 2, version B) which had similar syntax. For a given subject, two side-characters were randomly selected to be CN, and two side-characters were randomly selected to be UN. If a side-character was selected to be CN, one of the two possible CN event pairs was selected (e.g. Sandra Events 1 and 2, version A). If a side-character was selected to be UN, the two events were drawn from different possible CN event pairs, such that they belonged to unrelated narratives (e.g. Sandra Event 1, version A, and Event 2, version B). For instance, in one CN version for Sandra, she calls to discuss someone she is dating in Event 1, and the aftermath of that date occurs in Event 2. In one UN version for Sandra, she calls to discuss someone she is dating in Event 1, and she is seen hiding from art sponsors in Event 2.

This approach resulted in 32 possible arrangements of CN and UN events, which were pseudo-randomly assigned across subjects. For most subjects, 1 of 32 versions was randomly assigned by a seeded random number generator in MATLAB. After two-thirds of recruited subjects were scanned, it was determined that some versions had been under-sampled, therefore we generated a randomly-selected subset of the under-sampled versions to determine which version was assigned to the remaining subjects.

### Behavioral Tasks

For the fMRI study, on Day 1, subjects first completed consent forms, demographics questionnaires, and MRI screening. Prior to entering the scanning facility, participants completed brief familiarization tasks that were aimed at facilitating their attention to the story presentation task. Then, structural and functional imaging was performed, including the story presentation task. On Day 2, additional structural and functional imaging was performed, including the recall task. Familiarization, story presentation, and recall tasks were administered in MATLAB (https://www.mathworks.com/products/matlab.html) using Psychophysics Toolbox 3 (https://www.psychtoolbox.org). For event boundary ratings, an independent subject sample completed a one-day experiment in which they provided annotations of event boundaries.

#### Familiarization tasks (Day 1)

As described previously (Cohn-Sheehy et al., 2021), three brief tasks were administered to familiarize participants with character names and relationships, to orient participants to the upcoming stories, and to increase the likelihood that character names and relationships could be used as successful recall cues. Participants were first presented with character names, then character relationships, and then a test on character relationships (feedback provided). To encourage familiarity, these tasks incorporated face pictures for each character that were selected for high memorability (Bainbridge et al., 2013).

#### Story presentation (Day 1)

Stories were presented aurally using MRI-compatible Sensimetrics model S14 earbuds for binaural audio (https://www.sens.com/products/model-s14/). Stories 1-4 were, respectively, presented in scanning runs 1-4, and were presented again in the same order (Stories 1-4 in runs 5-8; **Fig. 1A**). This repetition of the stories was designed to increase the number of available trials and timepoints for pattern similarity analyses, which may benefit from large numbers of available trials. Prior to the task, participants were instructed verbally and onscreen that they would hear fictional clips involving characters from the familiarization tasks, and that they should devote their full attention and imagination to the clips as if they were listening to a book they enjoyed. Furthermore, they would later be asked to remember these stories in detail. Each story run began with 7 empty scanning timepoints, after which stories were presented binaurally through over-ear headphones, followed by 7 additional empty scanning timepoints. Only a white fixation cross was present onscreen during each story.

#### Recall task (Day 2)

Prior to performing the scanned recall task, participants were instructed that, after seeing the name of each character, they were to recall everything they could remember involving the particular character, from all stories, in as much detail as possible (Cohn-Sheehy et al., 2021). They were encouraged to attempt to recall as many details as possible for at least five minutes, and, if they were able to remember additional information, they were allowed to continue beyond five minutes. We did not instruct participants to recall events in any particular order, nor did we instruct participants to integrate information between events. Within one extended scanning run (**Fig. 1C**), participants were cued with the four CN and UN side-characters, in a randomized order. Each cue was presented onscreen, and included their relations to main protagonists, in order to encourage recall of both Events 1 and 2 (e.g. “Melvin Doyle (Charles Bort’s neighbor, Karen Joyce’s friend)”). After cueing CN and UN characters, cues were shown for the two main protagonists, in a randomized order. Each cue was preceded and followed by 7 empty scanning timepoints, with a white fixation cross onscreen. Spoken recall was recorded using an Optoacoustics FOMRI-III MRI-compatible microphone (http://www.optoacoustics.com/medical/fomri-iii).

#### Event boundary ratings

An independent sample completed a version of the Story Presentation task at a desktop computer (i.e. without brain imaging), with an additional instruction. Participants were presented with one of two versions of the story, and during story listening, were asked to press the spacebar whenever they perceived that one event had ended, and another event had begun.

### Recall scoring

Free recall performance was scored using a procedure developed in our previous behavioral experiments with a similar paradigm (Cohn-Sheehy et al., 2021), and which adapted a well-characterized scoring method from the Autobiographical Memory Interview (Levine et al., 2002). Recall data were scored in a blinded fashion by segmenting each participant’s typed recall into meaningful detail units, and then determining how many of these details could be verified within specific story events (**Data S2-3**). Briefly, each recall transcript was segmented into the smallest meaningful unit possible (“details”), and details were assigned labels that describe their content. We primarily sought to label details that could be verified within the story text (“verifiable details”). Verifiable details are details that specifically refer to events that center on the cued character (i.e. neither inferences about cued events, nor “external details” about other characters or events), which are recalled with some degree of certainty (e.g. not preceded by “I think” or “Maybe”), and which do not merely restate recall cues or other previously recalled details for a given cue.

We were also interested to discern recall performance for each CN or UN event (Events 1 or 2). For each CN or UN cue, verifiable details that referred to one particular event were labeled as “Event 1” or “Event 2.” If a verifiable detail could have originated from either Event 1 or Event 2, it was scored as “Either”; these details were rare (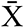 =0.06 details per cue, SD=0.24 details/cue, max=1.5 details/cue). Finally, if a verifiable detail merged information from both Event 1 and Event 2, it was marked as “Integrated”; these details were both rare and only observed for CN events (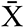 =0.14 details/cue, SD=0.56 details/cue, max=4.5 details/cue). To ensure that analyzed details were event-specific, Either and Integrated details were excluded from all reported analyses.

Audio transcription (AG, EM, JD, MD) and recall scoring (EM, JD, MD) was performed by individuals who were blinded to the experimental hypotheses and coherence conditions. Each participant’s recall transcript was first segmented by two raters, who were required to agree on the particular start and end times for each segment. Each rater subsequently scored the recall segments for verifiable details. Interrater reliability (IRR) for CN and UN events was high (mean Pearson’s *r*=0.95), and took into account how many verifiable details were scored per event label (1, 2, Either, Integrated), per character, and per participant. All reported comparisons between imaging and recalled details are based on counts of verifiable details for CN or UN events, averaged across raters.

### MRI acquisition

Scans were acquired at the UC Davis Imaging Research Center on a Siemens Skyra 3T scanner with a 32-channel head coil. Functional magnetic resonance imaging (fMRI) was performed using multi-band gradient-echo echo planar imaging (TR=1220 ms, TE=24ms, FA=67, MB Factor=2, FOV=19.2cm, matrix: 64 × 64, voxel size: 3.0mm isotropic, 38 slices). In order to isolate functional activity within particular structures of the brain, we also collected high-resolution structural imaging using a T1-weighted magnetization prepared rapid acquisition gradient echo (MPRAGE) pulse sequence (1 mm^3^ isotropic voxels). In order to ensure proper alignment between functional images between encoding and recall, we collected structural scans on both days.

On both Day 1 and Day 2, we first collected a brief localizer, and then the MPRAGE. We then adjusted the field of view for subsequent functional imaging to prioritize full inclusion of the temporal lobes (i.e. excluding some dorsal cortex, if necessary) and to align to the AC-PC axis. Initial shimming was then performed during a brief, task-free functional scan, after which we checked for artifacts (e.g. ghosting, aliasing) and determined whether the field of view needed to be adjusted (if so, this scan was repeated, with re-shimming). These preliminary scans were followed by functional imaging—story presentation scans on Day 1, and recall scans on Day 2. As time permitted, we additionally collected task-free functional imaging (Day 1), diffusion tensor imaging (Day 1 or Day 2), or additional task-concurrent imaging (Day 2, following the recall task). These additional scans are beyond the scope of the current experiment, and will be reported elsewhere.

### MRI preprocessing

Images were processed in SPM12 (https://www.fil.ion.ucl.ac.uk/spm/software/spm12/), including slice-time correction and realignment. After realignment, we checked for data quality using the MemoLab QA Toolbox (https://github.com/memobc/memolab-fmri-qa), which identifies excess motion and suspect timepoints or voxels within scanning runs using base functions in MATLAB, SPM, and ArtRepair (https://www.nitrc.org/projects/art_repair/), as well as detailed plots of signal intensity across all brain voxels and timepoints for each scanning run (Power, 2017). Furthermore, the QA Toolbox uses realignment parameters to create 6-directional motion regressors for each scan, which we implemented in subsequent models.

We excluded scanning runs if they exceeded 3mm motion in any direction, or if >20% of timepoints were identified as suspect based on framewise displacement >0.5mm or large deviations in signal intensity. We then merged realigned images into 4-dimensional files and used “film mode” in FslView (https://fsl.fmrib.ox.ac.uk/fsl/fslwiki/FslView/UserGuide) to inspect whether realignment failed – that is, whether there were any visible “jumps” between volumes after realignment. As previously mentioned, these approaches led us to exclude some subjects from participation in Day 2 (see Participants, above). When checking data quality for recall scans, we identified poor realignment due to large framewise displacement in two subjects. We attempted to salvage all data that preceded these moments of framewise displacement, by excluding subsequent volumes and re-performing realignment. This approach enabled us to salvage most of the recall scan for one subject (including all CN and UN recall), however, realignment still failed for the other subject. As such, one out of the twenty-five subjects could not be incorporated into encoding-retrieval similarity analyses.

Functional EPIs were co-registered with structural MPRAGE scans from the same day (Day 1 or Day 2). In order to compare functional volumes across days, we aligned images from Day 1 and Day 2, using the following approach. We co-registered Day 2 MPRAGEs to Day 1 MPRAGEs, and we applied these co-registration parameters to the realigned Day 2 images. This enabled the Day 2 functional images to directly overlap with Day 1 functional images. Additionally, we performed tissue segmentation on coregistered Day 1 MPRAGE scans, which we used to create brainmasks out of grey matter, white matter, and cerebrospinal fluid segmentations. For multi-voxel fMRI analyses (see below), these brainmasks were used to mask out extraneous voxels from first-level models of encoding and retrieval.

Furthermore, this enabled us to use the same structurally-defined regions of interest (ROIs) for both encoding and recall. We used a published probabilistic atlas of the medial temporal lobe (https://neurovault.org/collections/3731/) (Ritchey et al., 2015) to define the hippocampus, perirhinal cortex, and parahippocampal cortex, within each subject’s structural MPRAGE, by reverse-transforming these ROIs from template space (in MNI space) to native space (per subject) using Advanced Normalization Tools (ANTs; http://stnava.github.io/ANTs/). Each ROI was visually inspected against the native space scan. Because this parcellation separately defines the head, body, and tail of the hippocampus, we obtained whole-hippocampus ROIs by merging the head, body, and tail into one ROI on each side of the brain (left and right). As previously described (Reagh et al., 2020), for exploratory analyses of right hippocampal subregions (**Fig. S4**), we merged the right hippocampal body and tail ROIs to create a right posterior hippocampus ROI, and used the right hippocampal head as the right anterior hippocampus ROI; these ROIs had similar counts of voxels. Finally, other cortical ROIs which were implemented in post-hoc analyses (**Figs. S6-7**) were derived from individual subject parcellations in Freesurfer (https://surfer.nmr.mgh.harvard.edu/), using the Destrieux atlas (Destrieux et al., 2010). ROIs were then co-registered and resliced to functional images.

### Univariate measures of boundary-evoked activation

#### Boundary vs. Mid-Event timepoints

For univariate analysis of fMRI activation at event boundaries, structurally aligned BOLD data were smoothed with a 6mm-FWHM smoothing kernel and filtered with a 128s high-pass temporal filter prior to analysis. We modeled hippocampal activation at event boundaries using subject-level generalized linear models (GLMs) in SPM12. In order to control movement related confounds, all models include 6 rigid-body motion regressors (3 translational, 3 rotational). In line with published analyses of event boundary activation (Ben-Yakov & Henson, 2018), these models incorporated finite impulse response functions (FIRs) to model activation at event boundaries in a timepoint-by-timepoint fashion. The FIR approach enabled us to not only replicate the previous finding that hippocampal activation increases at event boundaries. It also enabled us to determine the lag between the onset of an event and the fMRI response to an event boundary. Furthermore, this enabled us to determine how long the hippocampus exhibits activation at event boundaries, such that we could use this epoch to constrain subsequent multivoxel fMRI analyses (modeled separately).

Event boundaries were defined *a priori*—they were intrinsic to the stimulus design (starts and ends of 40s events; **Fig. 1A**). Within the GLMs, we modeled activation at event boundaries (Boundary-Adjacent timepoints) versus Mid-Event timepoints (20s into each event), which did not contain any shifts in information that were expected to trigger event boundaries (see story text, **Data S1**). Importantly, we did not distinguish between specific types of events in this model (e.g. CN, UN, main plot). FIRs were modeled from -2 to +16 TRs around each timepoint—we selected this window in order to model all available event timepoints with minimal overlap between Boundary-Adjacent and Mid-Event windows. As such, the GLM resulted in beta images for each regressor, for each subject, which we then masked using the left and right hippocampus ROIs, averaging betas within each ROI, for each condition and each timepoint (e.g. Boundary-Adjacent TR +3, Mid-Event TR +14, etc.).

These averaged betas were then imported into RStudio for visualization and statistical analysis. Analyses of variance were performed using the Afex package in R (https://github.com/singmann/afex), including Greenhouse-Geisser corrections for any non-sphericity. For bilateral hippocampus, permutation tests were performed using Permuco (https://cran.r-project.org/web/packages/permuco/), testing for main effects at every timepoint using 100,000 permutations and a p<0.05 threshold for selecting clusters of significant timepoints, and then corrected for significance based on the mass of each cluster (Maris & Oostenveld, 2007). For post-hoc analyses of cortical networks (**Figs. S6-7**), the cluster threshold was adjusted for the number of models performed (3 models, therefore p<0.017). This approach enabled us to account for interdependency between adjacent timepoints, in line with well-documented approaches for time-series analysis of neuroimaging data (Cohen, 2014).

#### Activation during CN and UN events

To supplement the multivoxel fMRI analyses described below, we separately quantified univariate activation for CN and UN events, and incorporated this measure (“overall boundary activation”) as a fixed effect in mixed models of hippocampal pattern similarity during memory encoding (see Multi-level mixed effects models, below). In contrast to the univariate analysis reported above, this was performed using beta volumes which were initially modeled for multivoxel fMRI analysis (see Single trial models, below), and then smoothed with a 6mm-FWHM smoothing kernel for use in univariate analysis. We then extracted beta values during the Boundary epoch, and averaged these beta values within left and right hippocampus ROIs.

### Multivoxel fMRI analysis

#### Single trial models

We isolated the unique patterns of activity evoked by each CN or UN event during encoding, and each CN or UN cue during recall, using single-trial modeling (LS-S approach; Mumford et al., 2012; Turner et al., 2012). Specifically, we used finite impulse response functions (FIRs) to extract timepoint-by-timepoint activity in each voxel, for each modeled event for a given subject. These models were performed on unsmoothed data, in order to optimally characterize spatially-distributed patterns of activity (Kriegeskorte et al., 2006). For each story event (CN or UN), we modeled each timepoint (-2 TRs to + 45 TRs around the event onset) with a separate regressor, and used additional regressors to model each timepoint for all other story events within a given scanning run. This approach enabled the activity within each timepoint of a given event to be distinguished from activity from corresponding timepoints for all other events within a scanning run. The -2 to +45 TR window was designated to include extra available timepoints around the starts and ends of an event in order to: (1) differentiate activity corresponding with the event of interest from activity corresponding to adjacent events; and (2) capture the hemodynamic lag in the BOLD response around event boundaries (seen in the Univariate analysis of boundary-evoked activation, above).

For recall, we used a similar approach to model activity at each timepoint during a specific character cue, however, the number of available timepoints varied from cue to cue. For each recall cue, we therefore modeled timepoint regressors from -2 TRs before the onset of a recall cue to 7 TRs after the offset of that recall cue (corresponding with inter-cue times during scanning), and an equivalent number of regressors was designated to model activity from any other recall cue in the extended scanning run. For all models, we incorporated a 128s high-pass filter as well as 6 rigid-body motion regressors (3 translational, 3 rotational). This approach enabled us to distinguish patterns of activity that were evoked by each recall cue.

#### Representational similarity analysis

Once we isolated the unique patterns of activity associated with each story event and recall cue, we sought to test our hypotheses by comparing these patterns of activity across events. This type of analysis is referred to as Representational Similarity Analysis (RSA) (Kriegeskorte et al., 2008). In brief, RSA involves extracting the pattern of activity that is spread across a group of voxels in a given part of the brain (a “voxel pattern”), and computing correlations between voxel patterns that are evoked by different trials in an experiment. If voxel patterns are more highly correlated between trials in one experimental condition versus another, this provides evidence to suggest that whichever information distinguishes between those conditions is supported by that particular area of the brain.

To this end, we adapted several functions from the RSA Toolbox (http://www.mrc-cbu.cam.ac.uk/methods-and-resources/toolboxes/) and developed custom code in MATLAB. Voxel patterns were extracted within each ROI for a given subject, for every modeled timepoint for every trial (story event or recall cue). For each subject, and within each ROI, we then constructed a representational similarity matrix (RSM), by correlating voxel patterns (Pearson’s r) from every modeled timepoint with every modeled timepoint.

To test our hypotheses about how activity patterns are shared between CN versus UN events, and between these events and recall, we selectively averaged correlations from the overall RSM. For analyses of encoding, we selected all correlations between timepoints from two events involving the same CN or UN character (e.g. Beatrice Event 1 vs Beatrice Event 2). For encoding-retrieval similarity, we selected all correlations between timepoints from a CN or UN encoding event and timepoints corresponding to the recall cue for the same CN or UN character (e.g. Beatrice Event 1 vs Beatrice recall cue). Importantly, all of these correlations were computed across scanning runs (at least two scans apart), such that any effects of temporal autocorrelation would be minimized. After selecting these specific sets of correlations, we averaged them to derive one condition-specific, timepoint-by-timepoint correlation matrix for each ROI, for each subject. As described below, within each condition, we then averaged pattern similarity values (mean Pearson’s r) within empirically-defined epochs (e.g. Boundary epoch).

#### Across-event similarity epochs

To test our specific hypotheses about representational similarity involving event boundaries, we computed selective averages of each condition-specific RSA matrix (described above) based on empirically-derived Boundary and Post-Boundary epochs, which were then imported into RStudio for statistical analysis. Within each across-event similarity matrix, for each subject and ROI, we selectively averaged timepoint-by-timepoint correlations between events (**Fig. 2**): Boundary-Boundary similarity, between TRs +5 to +11 from one event, and the same TRs from the other event; Boundary-Post-Boundary similarity, between TRs +5 to +11 from one event, and TRs +12 to +37 from the other event; and Post-Boundary-Post-Boundary similarity, between TRs +12 to +37 from one event, and the same TRs from the other event. These three measures of epoch-by-epoch pattern similarity constitute the three levels of the “Epoch” factor within mixed models of across-event pattern similarity (see Multi-level mixed effects models, below).

#### Follow-up timepoint-by-timepoint analysis

As shown in **Fig. S2**, we also analyzed timepoint-by-timepoint pattern similarity between pairs of events at encoding, to follow-up on epoch-based analyses (i.e. to see which timepoints had CN>UN similarity). To assess statistical significance for narrative coherence effects, and to correct for multiple comparisons, we used cluster-based permutation tests (Maris & Oostenveld, 2007) with 10,000 permutations, with a cluster defining threshold of p=0.05 (one-tailed, to test for CN>UN similarity) and a cluster mass threshold of p=0.05. Each pixel of a statistical comparison (t-value) was converted into a Z value by normalizing it to the mean and standard error generated from our permutation distributions. Above-threshold (Z>1.96) pixels and clusters are plotted in **Fig. S2C-D**.

#### Encoding-retrieval similarity

Within the right hippocampus, for each encoding event (Event 1 or 2), we selectively averaged timepoint-by-timepoint correlations between timepoints spanning both Boundary and Post-Boundary epochs at encoding (i.e. the whole event), and any timepoints during recall of the same CN or UN character (**Fig. 3**). This yielded two measures of encoding-retrieval similarity: Event 1 Reinstatement and Event 2 Reinstatement. These measures were specified by a factor of Event Number [Event 1 Reinstatement, Event 2 Reinstatement] within mixed models (see Multi-level mixed effects models, below). For ancillary analyses, we also selectively averaged correlations between recall and the Boundary epoch alone (**Fig. S5**).

### Textual similarity analysis

We modeled textual similarity between sentences from different story events using the freely available, “transformer” version of the Universal Sentence Encoder (USE), a text embedding model designed to convert text into numerical vectors (Cer et al., 2018; see also, https://ai.googleblog.com/2018/05/advances-in-semantic-textual-similarity.html). Briefly, the USE uses pre-weighted layers, pre-trained on an expansive textual database, to transform inputted sentences into 512-dimensional embedding vectors that account for the combination of words, and the respective positions of words, within each sentence. Then, cosine similarity is calculated between each sentence vector, yielding pairwise measures of textual similarity for all inputted sentences. As such, USE similarity serves as a proxy of word- and sentence-level semantic relatedness for text.

For each subject, we created a textual-similarity matrix for cosine similarity between all sentences from all stories. In order to compare textual similarity with fMRI voxel pattern similarity, we used custom MATLAB code to align the whole USE similarity matrix with the whole RSM for each subject. Because all sentences were of equivalent lengths (5s each), we were able to align the time-course of sentences with the time-course of fMRI scans (1.22s for each timepoint). However, in order to align the 8-sentence window with the 40s window for each event, it was necessary to only include fMRI timepoints that corresponded with that 40s window. Therefore, we excluded any timepoints that did not correspond with Boundary or Post-Boundary event epochs (i.e. including only TRs +5 to +37, 40s total). The whole USE similarity matrix was upsampled and interpolated from a 5s/sentence timescale to match the 1.22s/TR timescale of the whole RSM. Finally, USE similarity was selectively averaged within each condition of interest (CN vs UN) and time-window (Boundary-Boundary, Boundary-Post-Boundary, Post-Boundary-Post-Boundary). These averaged USE similarity values were subsequently incorporated into mixed-effects models (see Statistical Analysis, Formula 2, below).

### Statistical Analysis

Statistical tests were performed in R using standard functions t-tests, as well as Afex (https://github.com/singmann/afex) for mixed models and ANOVAs, and Permuco (https://cran.r-project.org/web/packages/permuco/) for permutation tests. For ANOVAs, Greenhouse-Geisser corrections were implemented for any non-sphericity. For visualization (**Figs. S1, S7**), within-subjects standard error was calculated using Rmisc (https://cran.r-project.org/web/packages/Rmisc/index.html), by normalizing data to remove between-subject variability, and computing variance from this normalized data (Morey, 2008).

#### Multi-level mixed effects models

Mixed model formulas are reported within the Results text. Fixed effects specifications were based on experimental hypotheses, random effects structures which accounted for effects of individual subjects (“Subject”) and the content of particular events (“Event Content”). For models of across-event similarity at encoding, effects of Event Content (i.e. “item” effects) were operationalized as the particular pair of events which were presented for a particular CN or UN character (e.g. Beatrice Events 1A+2B, Melvin Events 1B+2B). For models of encoding-retrieval similarity or recalled details, Event Content was operationalized as which particular event was being recalled (e.g. Beatrice Event 1A, Beatrice Event 2B, Melvin Event 1B, or Melvin Event 2B). Random effects structures were specified by first attempting to fit a maximal random effects structure justified by the experimental design (including random slopes) (Barr et al., 2013), followed by systematically pruning the random effects structure until the model converged without a singular solution (i.e. without overfitting; Singmann & Kellen, 2019). Because models which included random slopes either did not converge or reached a singular solution, reported models only included intercepts for random effects. This approach resulted in the following model formulas:

Formula 1 (**Fig. 2B**): Pattern similarity ∼ Narrative Coherence * Epoch + Overall Boundary Activation + (1 | Subject) + (1 | Event Content)

Formula 2: Pattern similarity ∼ Narrative Coherence * Epoch + Overall Boundary Activation + USE similarity + (1 | Subject) + (1 | Event Content)

Formula 3 (**Fig. 3B**): Encoding-retrieval similarity ∼ Narrative Coherence * Event Number + Recall Duration + (1 | Subject) + (1 | Event Content)

Formula 4 (**Fig. 3C**): Recalled details ∼ Event 2 Reinstatement * Narrative Coherence * Event Recalled + Recall duration + (1 | Subject) + (1 | Event Content)

Model fits were quantified using the Akaike Information Criterion (AIC), and significance for fixed effects was calculated using the Kenward-Roger approximation for degrees of freedom, which is useful for unbalanced mixed designs (Singmann & Kellen, 2019). To follow up on significant fixed effects and interactions, we used the emmeans package (https://cran.r-project.org/web/packages/emmeans/index.html) to conduct pairwise comparisons and visualization of factors (emmeans; **Figs. 2-3, S3-6**) and interactions between factors and continuous variables (emtrends and emmip; **Fig. 3C**, **Fig. S5C**).

## Acknowledgments

We thank Alexander Garber, Elena Markantonakis, Mehar Durah, and June Dy for transcribing and scoring recall. We thank Jesse Doty, Sarah Morse, Costin Tanase, Dennis Thompson, and the Imaging Research Center for their technical contributions. We thank Halle Dimsdale-Zucker, Derek Huffman, Anna Leshinskaya, Walter Reilly, Andy Yonelinas, and the Dynamic Memory Lab (http://dml.ucdavis.edu) for consultation on experimental design and analysis, and James Antony and Cameron Riddell for feedback on the manuscript. This research was supported by a Multi-University Research Initiative grant N00014-17-1-2961 from the United States Office of Naval Research/Department of Defense (CR, JMZ), a Ruth L. Kirchstein National Research Service Award F30AG062053 from the National Institute on Aging (BICS), and a Floyd and Mary Schwall Medical Research Fellowship from the University of California, Davis (BICS). Any opinions, findings, conclusions, or recommendations expressed in this material are those of the authors and do not necessarily reflect the official views of the Office of Naval Research, U.S. Department of Defense, or National Institutes of Health.

---

**Fig. S1.**
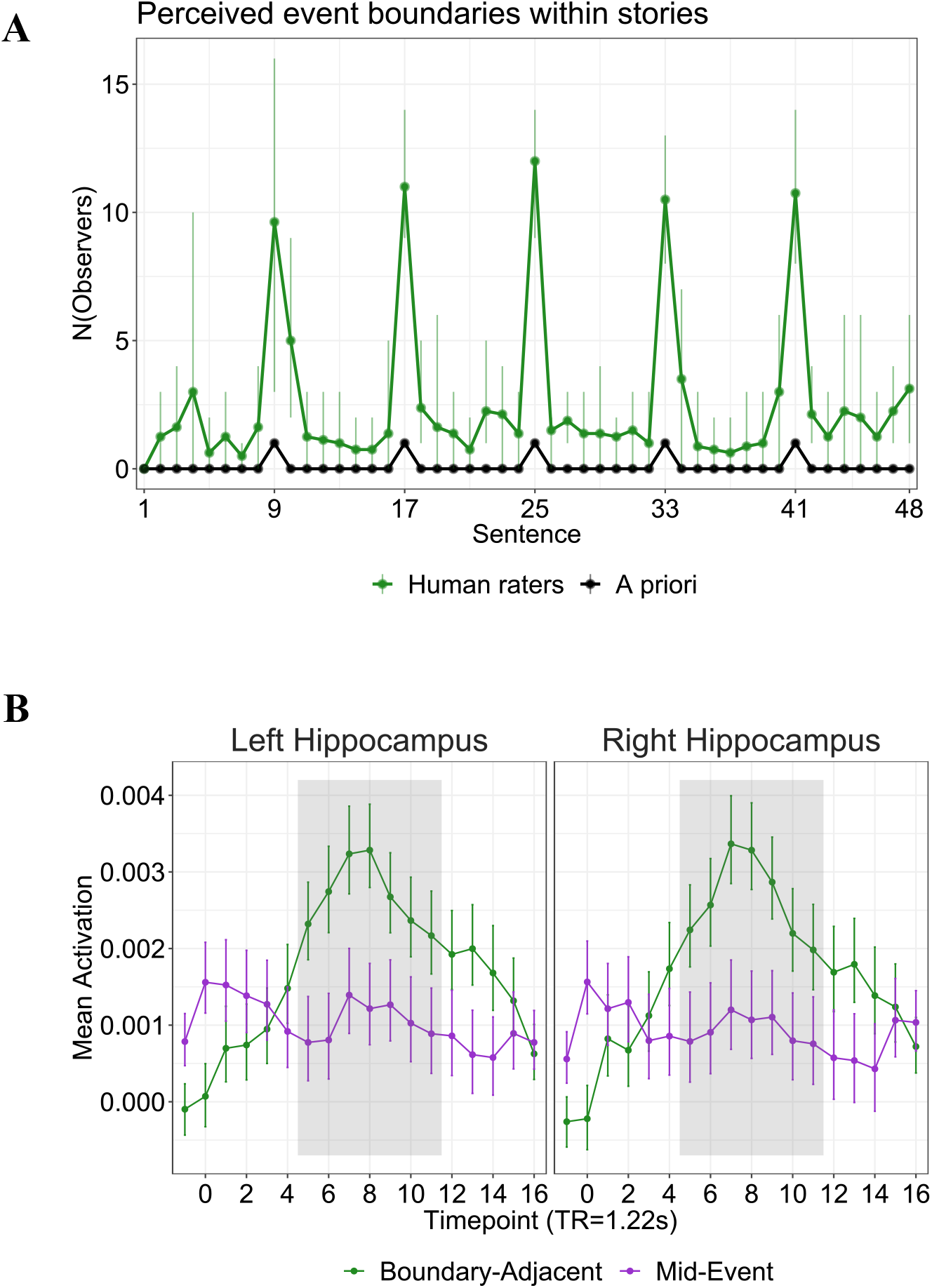
A priori event boundaries evoke boundary perception and increased hippocampal activation. Each story was written to include *a priori* event boundaries, such that it would naturally be segmented into six 40-second events (see Methods). **(A)** Perceived boundaries correspond with *a priori* boundaries: An independent sample (N=18, not scanned) annotated perceived event boundaries during story presentation. The total number of observers who perceived a boundary during each 5-second story sentence (N(Observors), green; dots/line = mean across stories, vertical lines = range) was compared with the locations of *a priori* event boundaries (black dots/line; not including story onset and offset). **(B)** Boundary-evoked hippocampal activation: BOLD activation time-courses were modeled in the left and right hippocampus, to estimate activity changes at Boundary-Adjacent (green; -2 to +16 TRs around an a priori event boundary) versus Mid-Event timepoints (purple; 20 seconds into each 40-second event). These activation time-courses were modeled using an analysis of variance (ANOVA) with three within-subjects factors: Condition [Boundary-Adjacent vs Mid-Event], Timepoint, and Hemisphere [left vs right hippocampus]. This analysis revealed significant main effects of Condition (*F*(1,24)=29.0, η_G_^2^=.007, p<.0001) and Timepoint (*F*(2.56, 61.37)=4.92, η_G_^2^=.02, p=.006) that were qualified by a significant Condition by Timepoint interaction (*F*(5.06, 121.53)=11.2, η_G_^2^=.02, p<.0001; for other effects, ps>.15). Following up on the significant Condition by Timepoint interaction, cluster-based permutation testing revealed a cluster of Boundary-Adjacent timepoints where activity was significantly higher than corresponding Mid-Event timepoints (grey shading; TRs +5 to +11; cluster-defining threshold is p<.05). We incorporated both the left and right hippocampus into this analysis because we did not observe any effects of Hemisphere in the preceding ANOVA. Means are represented by dots and lines, and error bars represent within-subjects estimates of standard error (see Methods for details).

**Fig. S2.**
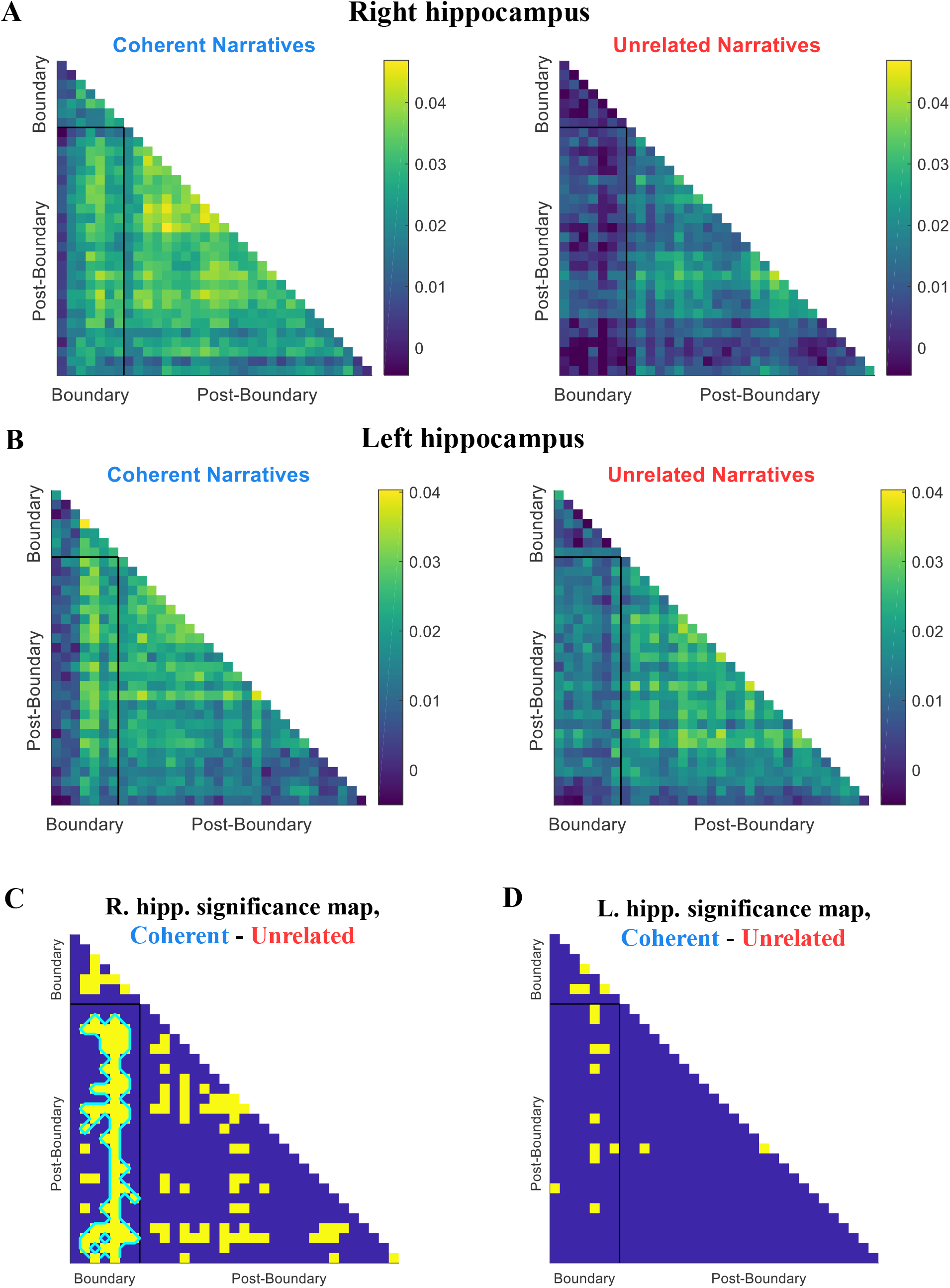
Timepoint-by-timepoint similarity in the right and left hippocampus. (A-B) Mean pattern similarity: Pattern similarity (Pearson’s R) between Event 1 and 2 was computed for Coherent Narrative and Unrelated Narrative events, between timepoints from the Boundary epoch (TRs 5-11) and Post-Boundary epoch (TRs 12-37). Here, pattern similarity within the right hippocampus (A) and left hippocampus (B) is averaged across subjects, within Narrative Coherence conditions, for each timepoint-by-timepoint correlation. (C-D) Significant timepoint-by-timepoint pattern similarity (see Methods): within the right hippocampus (C), several timepoint-by-timepoint correlations were significantly different between Coherent Narrative and Unrelated Narrative conditions (yellow, Z>1.96). This included a significant cluster of timepoint-by-timepoint similarity between the Boundary epoch of one event, and the Post-Boundary epoch of another event (outlined in cyan). These findings were not evident in the left hippocampus (D).

**Fig. S3.**
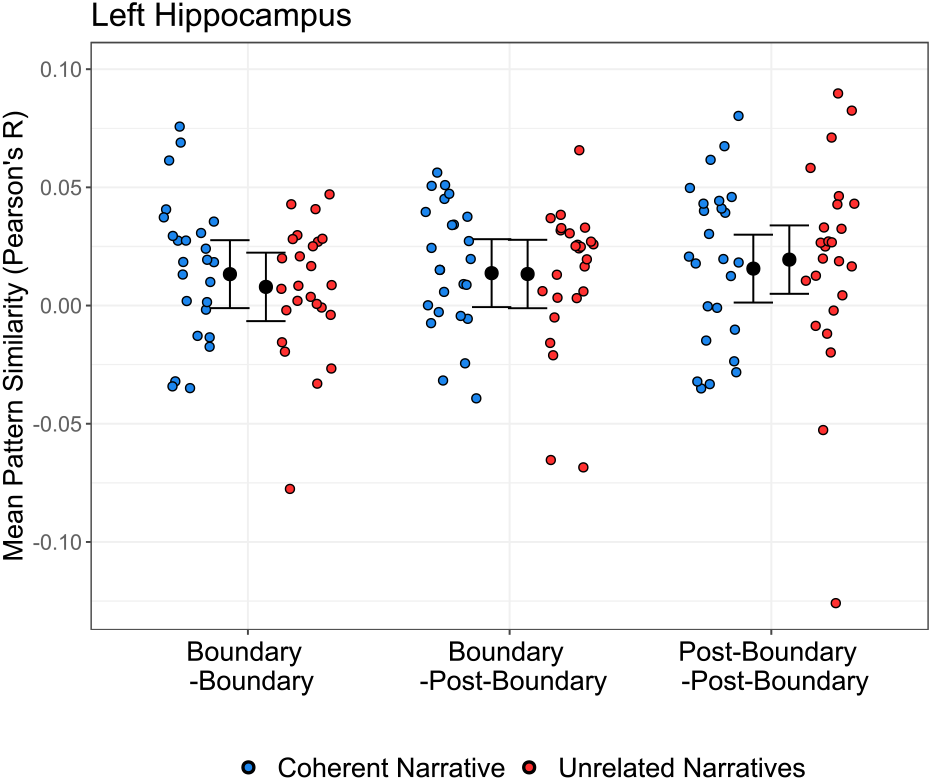
No effect of narrative coherence on left hippocampus pattern similarity. Left hippocampus pattern similarity between Event 1 and 2 epochs was computed for Coherent Narrative and Unrelated Narrative events. Pattern similarity for individual subjects (colored dots) is plotted on the basis of which epoch was being examined, and whether events could form Coherent Narratives or not (Unrelated Narratives). Mixed model estimates predicting pattern similarity are visualized as 95% confidence intervals (see text for model details). Key: Blue=Coherent Narratives, Red=Unrelated Narratives.

**Fig. S4.**
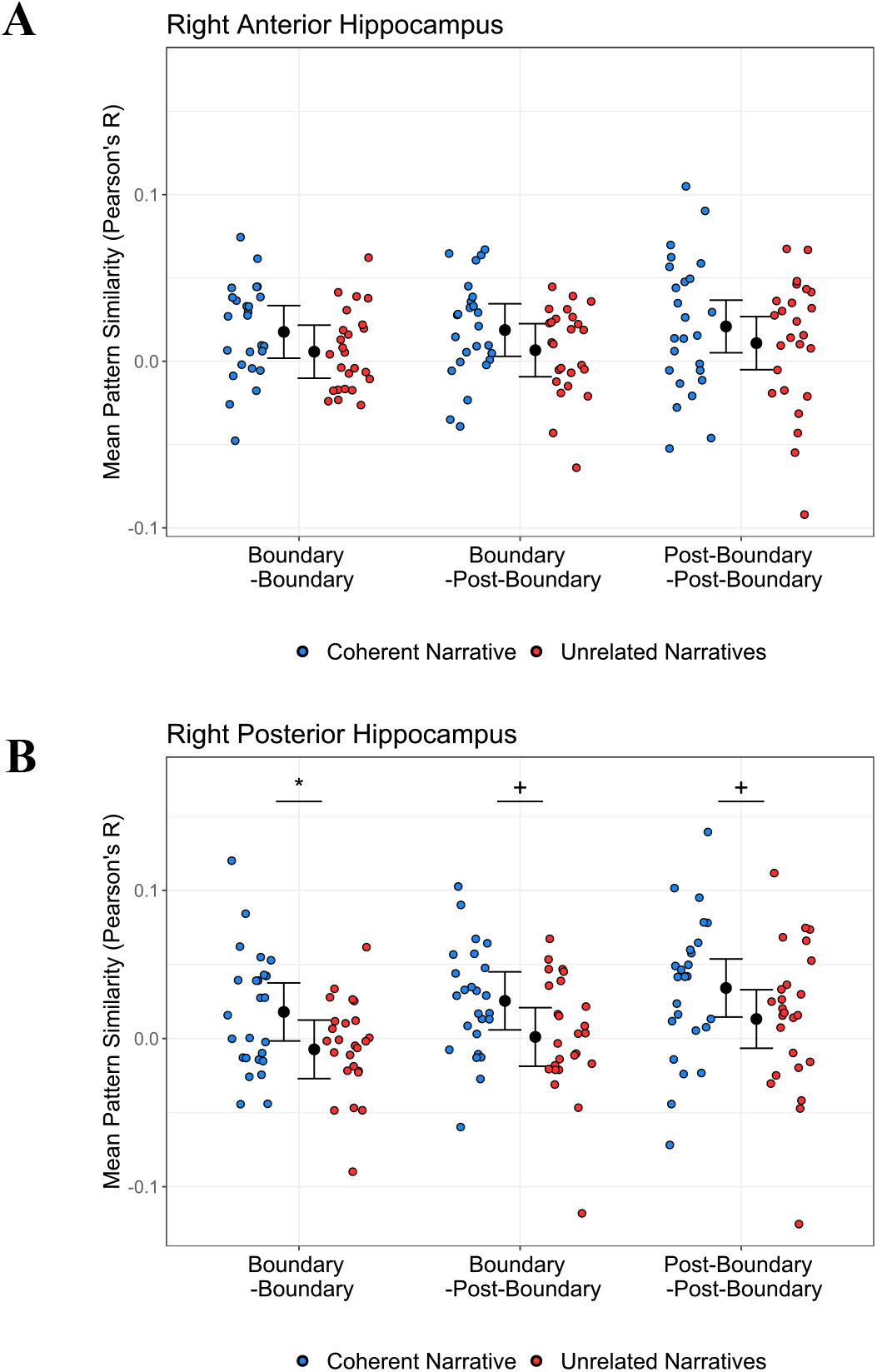
Exploratory pattern similarity analysis in right anterior and posterior hippocampus. Following up on findings in the right hippocampus (Figure 3), pattern similarity between Event 1 and 2 epochs was computed for Coherent Narrative and Unrelated Narrative events, within the right anterior **(A)** and posterior hippocampus **(B)**. Pattern similarity for individual subjects (colored dots) is plotted on the basis of which epoch was being examined, and whether events could form Coherent Narratives or not (Unrelated Narratives). Mixed model estimates predicting pattern similarity are visualized as 95% confidence intervals (similar to Figure 3); pairwise differences in right posterior hippocampus reflect a significant main effect of Narrative Coherence on pattern similarity (*F*(1,11.66)=6.85, p=0.023; model AIC=-868.56). This main effect was not significant within the right anterior hippocampus (*F*(1,11.73)=2.16, p=0.17; model AIC=-998.62). Key: Blue=Coherent Narratives, Red=Unrelated Narratives; *=p<.05, +=p<.10, uncorrected for multiple comparisons.

**Fig. S5.**
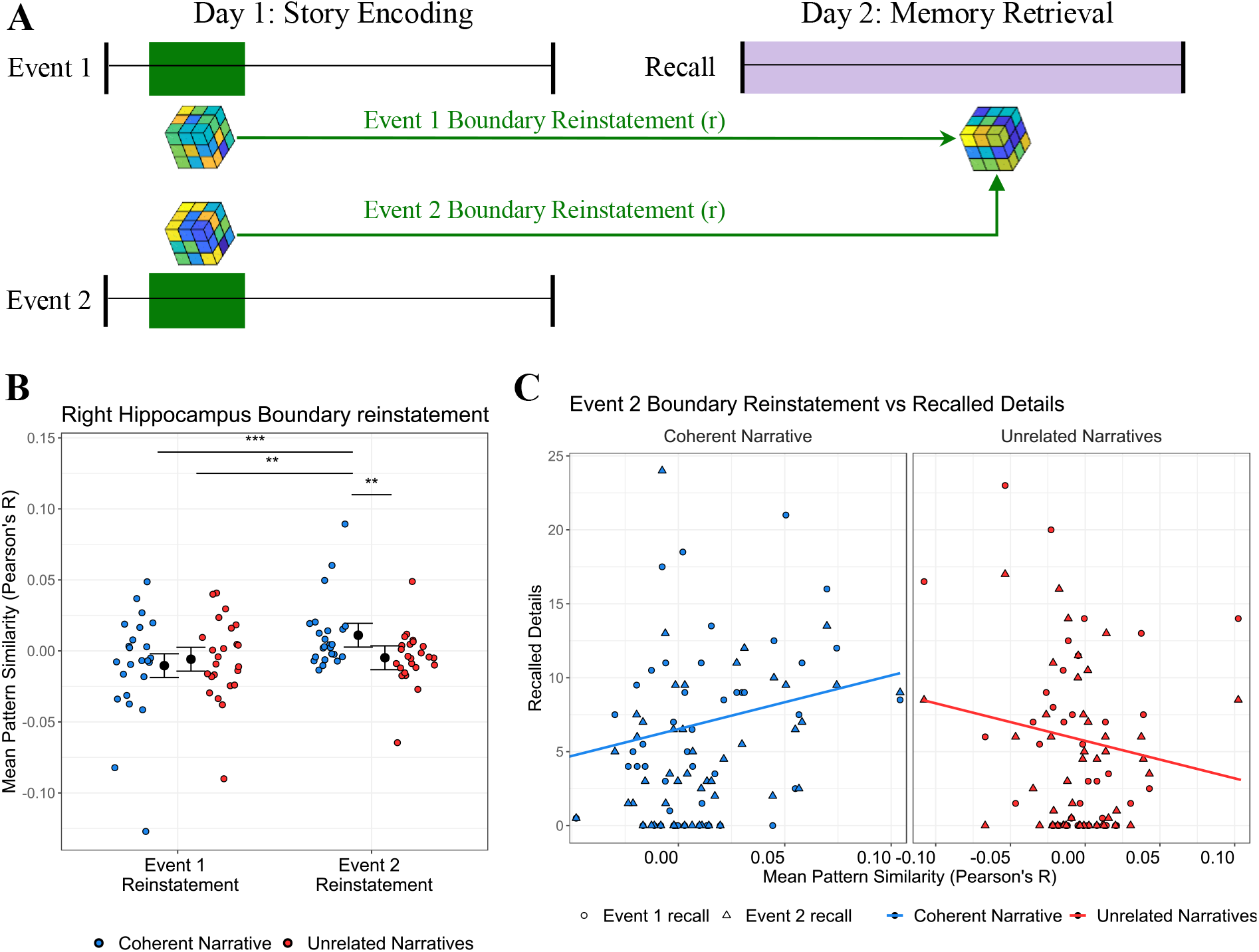
Reinstatement of boundary-evoked hippocampal activity from memory encoding supports recall of Coherent Narrative events. **(A)** Encoding-retrieval similarity: Pattern similarity (Pearson’s r) was calculated between boundary-evoked activity from each Event (Event 1 Boundary, Event 2 Boundary) involving a recurring character at encoding, and delayed recall of events involving that character. **(B)** Event 2 boundary pattern reinstatement is higher for Coherent Narrative recall: Encoding-retrieval similarity for each subject (colored dots) is plotted for each epoch of each event during recall. 95% confidence intervals represent mixed model estimates (AIC=-1427.5; interaction of Narrative Coherence by Event Number: *F*(1,343.00)=7.58, p=.006). **(C)** Event recall predicted by Event 2 boundary pattern reinstatement: Recalled details from Event 1 (circles) and Event 2 (triangles) are plotted for each subject, predicted by Event 2 boundary pattern reinstatement during recall. Colored lines represent mixed model estimates (AIC=1114.7; interaction of Event 2 Reinstatement by Narrative Coherence: *F*(1,160.15)=10.56, p=.001), which were significantly different (*t*(160)=3.25, p=.001, uncorrected for multiple comparisons). Key: Blue=Coherent Narrative, Red=Unrelated Narratives; **=p<.01, ***=p<.001, uncorrected for multiple comparisons.

**Fig. S6.**
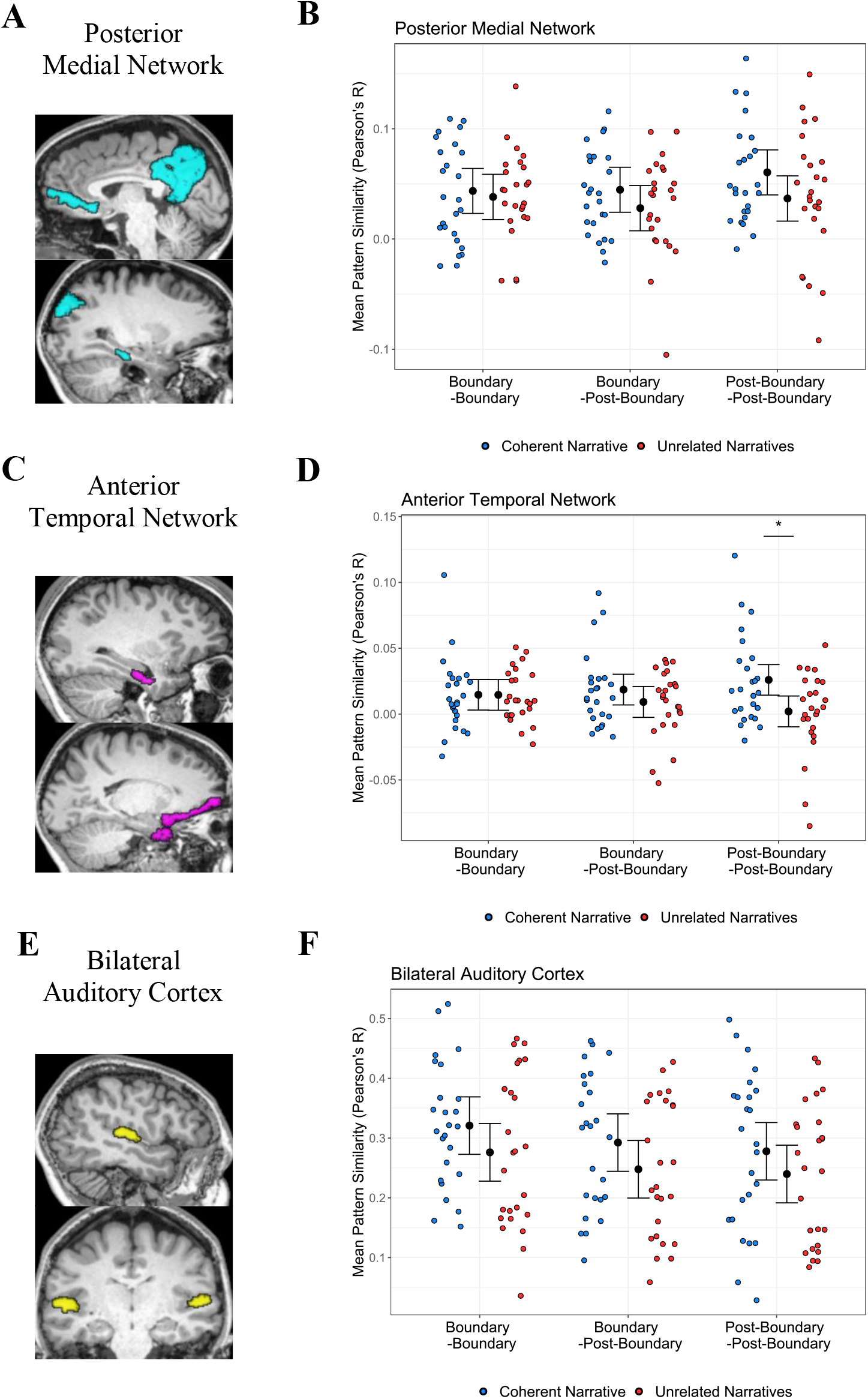
Post-hoc analysis of narrative coherence effects in cortical networks. **(A,C,E)** Regions of interest within three cortical networks are overlaid on a single-subject brain: the Posterior Medial Network (**A**: medial prefrontal cortex, posterior medial cortex, angular gyrus, parahippocampal cortex), Anterior Temporal Network (**C**: perirhinal cortex, orbitofrontal cortex, temporal pole), and Bilateral Auditory Cortex (**E:** transverse temporal gyrus). (**B,D,F**) Similar to Figure 3, Pearson’s correlations between Event 1 and 2 epochs were computed for Coherent Narrative and Unrelated Narrative events. Pattern similarity for individual subjects (colored dots) is plotted on the basis of which epoch was being examined, and whether events could form Coherent Narratives or not (Unrelated Narratives). 95% confidence intervals (black) represent the predictions of mixed models with the following formula: Pattern similarity ∼ Narrative Coherence * Epoch * Region of interest (ROI) + (1 | Subject) + (1 | Event Content). Pairwise comparisons following up on significant fixed effects were corrected for multiple comparisons using the Holm method. (**B**) Within the Posterior Medial Network (AIC=-4372.2), this model revealed significant effects of Epoch (*F*(2,2315.22)=3.38, p=.034) and ROI (*F*(7,2315.22) = 19.05, p<0.001; not shown here), however, the effect of Narrative Coherence was not significant (*F*(1,12.35)=2.20, p=0.16; for interactions, ps>.15). (**D**) Within the Anterior Temporal Network (AIC=-5159.5), there was a significant effect of ROI (*F*(5,1727.42)=3.04,p=.01; not shown here) and a significant interaction between Narrative Coherence and Epoch (*F*(2,1727.42)=6.95, p<.001; all other effects, p>.07). (**F**) Within bilateral auditory cortex (AIC=-608.1), there were significant effects of ROI (F(1,552.88)=18.61, p<0.001; not shown here), Epoch (*F*(2,552.88)=5.10, p=0.006), and Narrative Coherence (*F*(1,8.81)=13.21, p=0.006). Key: Blue=Coherent Narratives, Red=Unrelated Narratives, *=p<0.05.

**Fig. S7.**
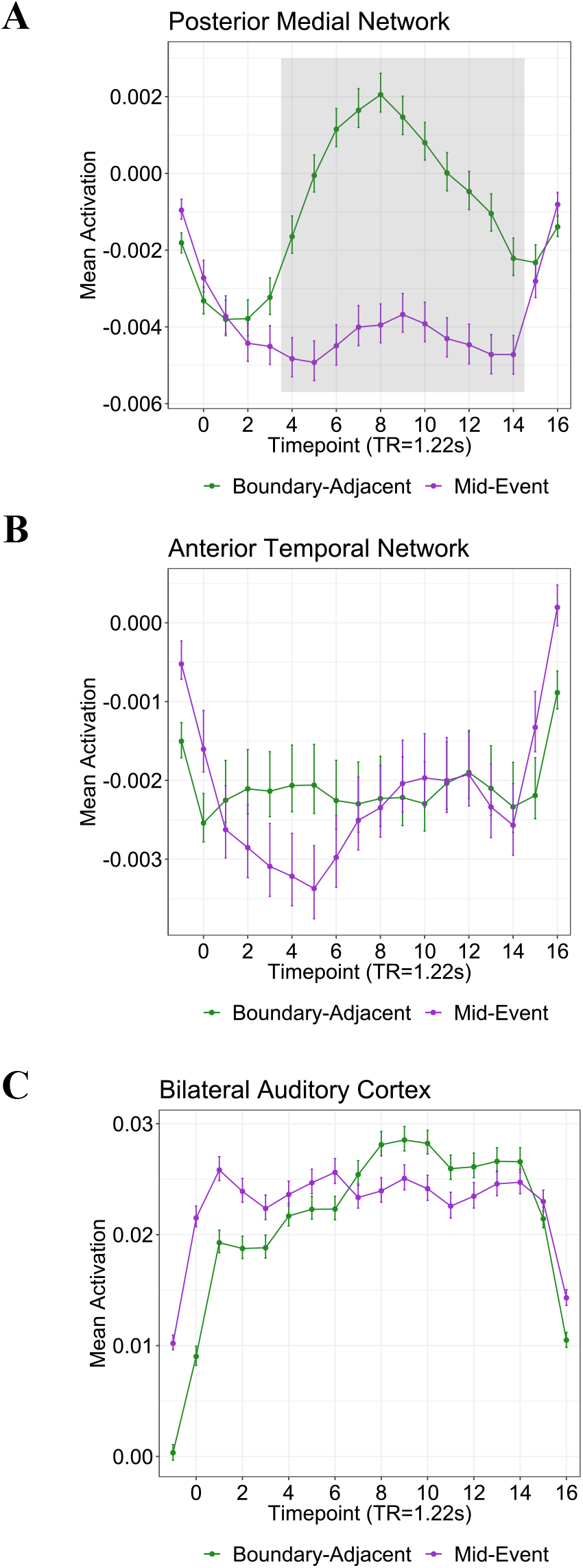
Post-hoc analysis of boundary-evoked activation in cortical networks. Similar to Figure 2B, Boundary-Adjacent activation (green) is compared with Mid-Event activation (purple) within the Posterior Medial Network **(A)**, Anterior Temporal Network **(B)**, and Bilateral Auditory Cortex **(C)**. Means are represented by dots and lines, and error bars represent within-subjects estimates of standard error (see Methods for details). Grey shading (in **A**) represents a significant cluster of boundary activation in the Posterior Medial Network (TRs +4 to +14; cluster-defining threshold corrected for multiple comparisons, p<0.017).

